# Engineering a niche supporting haematopoietic stem cell development using integrated single cell transcriptomics

**DOI:** 10.1101/2021.01.25.427999

**Authors:** Brandon Hadland, Barbara Varnum-Finney, Stacey Dozono, Tessa Dignum, Cynthia Nourigat-McKay, Dana L Jackson, Tomer Itkin, Jason M. Butler, Shahin Rafii, Cole Trapnell, Irwin D. Bernstein

## Abstract

Haematopoietic stem cells (HSCs) develop from haemogenic endothelium (HE) within embryonic arterial vessels such as the aorta of the aorta-gonad-mesonephros region (AGM). To identify the signals responsible for HSC formation, we used single cell RNA-sequencing to simultaneously analyze the transcriptional profiles of AGM-derived cells transitioning from HE to HSC, and AGM-derived endothelial cells which provide signals sufficient to support HSC maturation and self-renewal. Pseudotemporal ordering revealed dynamics of gene expression during the HE to HSC transition, identifying surface receptors specifically expressed on developing HSCs. Transcriptional profiles of niche endothelial cells enabled identification of corresponding ligands, including those signaling to Notch receptors, VLA-4 integrin, and CXCR4, which, when integrated in an engineered platform, were sufficient to support the generation of engrafting HSCs. These studies provide a transcriptional map of the signaling interactions necessary for the development of HSCs and advance the goal of engineering HSC for therapeutic applications.

## Introduction

One of the longstanding challenges in the field of haematopoiesis has been identifying the unique signal pathways and transcriptional programs that orchestrate the formation of long-term engrafting haematopoietic stem cells (HSCs). Complicating the study of HSC formation, embryonic haematopoiesis proceeds through sequential waves of lineage-restricted primitive, erythroid/myeloid, and lymphoid/myeloid progenitors prior to the emergence of the first HSCs^1^. During haematopoietic development, specialized endothelial-like precursor cells, termed haemogenic endothelium (HE), simultaneously downregulate endothelial-specific and activate haematopoietic-specific transcriptional programs to give rise to haematopoietic progenitors and HSCs. For functional long-term engrafting HSC to emerge, HE must acquire and maintain HSC-defining properties such as the ability to self-renew, home, and provide multilineage haematopoiesis, properties which distinguish rare HSCs from other embryonic haematopoietic progenitors. Currently, the combination of cell intrinsic and extrinsic signals required to support the acquisition of these HSC-defining properties during embryonic development of HE remains inadequately defined.

The emergence of HSCs occurs uniquely in the context of arterial vessels such as the aorta of the embryonic region referred to as the aorta-gonad-mesonephros (AGM)^2-4^. The arterial environment in which HSCs develop suggests cell intrinsic properties of “arterialized” HE and cell extrinsic signals from the arterial vascular niche may contribute to HSC fate. Supporting this concept, a number of studies have demonstrated the importance of arterial programs, such as those regulated by the Notch pathway, explicitly in definitive multilineage haematopoiesis and HSC formation^5-12^. This argues the need for further efforts to identify the precise signaling interactions between arterialized HE and the arterial vascular niche required for HSC formation.

Further complicating the study of HSCs is their scarcity relative to other progenitors arising simultaneously in the developing embryo. HSC activity, as measured by direct transplantation into adult immune-competent recipients, is first consistently detected at around embryonic day 11 (E11) in the AGM at very low frequency (∼1 HSC per AGM based on limit dilution transplantation)^13^. Haemogenic precursors capable of maturation to HSC by ex vivo stromal culture or neonatal transplantation have been detected as early as E9 and increase in numbers between E10 and E11 in the AGM, prior to migration to the fetal liver where further maturation and expansion of HSCs occurs^13-15^. These HSC precursors have been characterized by their sequential expression of initial haematopoietic surface markers, CD41 followed by CD45, within the larger population of cells expressing the endothelial marker VE-Cadherin, defining a series of VE-Cadherin^+^CD41^+^CD45^-^ pro-HSC/pre-HSC I and VE-Cadherin^+^CD45^+^ pre-HSC II that mature to functional, long-term engrafting VE-Cadherin^+/-^ CD45^+^ HSCs^16-18^. Although this phenotypic characterization of HSC precursors has been helpful in defining the sequence of HSC emergence, our understanding of the unique molecular properties of developing HSCs is limited by the fact that HSCs arise asynchronously from HE and remain far outnumbered by other haematopoietic progenitors with overlapping phenotypes differentiating simultaneously in intra-aortic haematopoietic clusters between E9 and E11^19^. This argues the need to study the unique properties of rare HSC precursors using single cell techniques.

Toward this goal, advances in single cell RNA sequencing (scRNA-seq) technology and complementary improvements in computational algorithms to explore large, complex single cell data sets have enabled transcriptome-wide analysis of rare stem cell populations and transitional states in development at unprecedented resolution^20^. Recent efforts have elegantly applied scRNA-seq to study the emergence of HSCs, providing new insights into the transcriptional properties of HSC precursors^21-24^. Building on these studies, we report here the application of single cell transcriptomic approaches to address the complex interactions between the arterial vascular niche and HE in regulating the specification and self-renewal of HSCs. We leveraged our previously reported model utilizing AGM-derived endothelial cell stroma (AGM-EC), which provides an instructive niche for the generation of engrafting HSCs from clonal embryonic precursors as early as E9^25-27^, to identify a cell surface phenotype (VE-Cadherin^+^CD61^+^EPCR^+^) enriching for HSC precursors across their developmental spectrum from HE. We then applied scRNA-seq to this enriched population to study the transcriptional changes associated with HSC formation in the vascular niche at single cell resolution, focusing analysis to genes encoding cell surface receptors and downstream signaling molecules. Complementary transcriptomic analysis of the AGM EC niche identified cognate ligands interacting with receptors on developing HSCs, which enabled us to engineer a stromal cell-independent niche that supported the generation of engrafting HSCs from embryonic haemogenic precursors in vitro. Altogether, these studies provide novel insight into the precise conditions necessary to recapitulate the development of HSCs from HE, advancing translational efforts for de novo HSC generation.

## RESULTS

### Single cell RNA-sequencing identifies the transcriptional signatures of AGM-derived niche endothelial cells supporting HSC emergence

Identifying the microenvironmental signals necessary for generation of functional HSCs requires analysis of niche cells capable of supporting the process of HSC specification and self-renewal. We previously demonstrated that Akt-activated AGM-derived EC (AGM-EC), which retain characteristics of their primary cells of origin including expression of Notch ligands, provide an instructive in vitro niche for the generation and expansion of HSCs from embryonic haemogenic precursors^25^. Most independently generated AGM-EC support the production of long-term, multilineage-engrafting HSCs following co-culture of AGM-derived VE-Cadherin^+^ haemogenic precursors isolated at E9-E10; however, we identified rare AGM-EC that are deficient in this capacity (Fig. 1a-b). To determine differences in transcriptional properties of HSC-supportive and non-supportive AGM-EC, we compared their genome-wide transcriptional profiles by scRNA-seq. Dimensionality reduction by UMAP and clustering of single cell transcriptomes revealed that the non-supportive AGM-EC cluster apart from three independent HSC-supportive AGM-EC in transcriptional space (Fig. 1c, Supplementary Fig. 1). We then examined genes differentially expressed between the HSC-supportive and non-supportive AGM-EC, identifying genes most specific to the HSC-supportive cluster, which we hypothesize to include genes encoding signaling ligands essential for HSC generation (Supplementary Table 2). Gene ontology analysis of this gene set suggests a prominent role in HSC-supportive AGM-EC for genes involved in regulation of cell adhesion, integrin interactions, signaling receptor/growth factor binding, and cell death/apoptosis (Fig. 1d, Supplementary Table 3). Amongst the top significantly differentially expressed genes were transcription factors implicated in arterial EC and HSC fates, such as *Sox17*^7, 28-30^, genes encoding secreted signaling ligands, such as the chemokine *Cxcl12*, and genes encoding cell adhesion and extracellular matrix proteins, particularly those interacting with integrins, such as *Icam1, Vcam1, Fn1* (Fibronectin), and *Tgm2* (Tissue transglutaminase) (Fig. 1e, Supplementary Table 2). These results identify candidate signals that may support the maturation and self-renewal of HSCs from haemogenic precursors in the AGM-EC niche, as well as transcription factors that may regulate the differential expression of niche signals between HSC-supportive and non-supportive AGM-EC.

**Figure 1.**
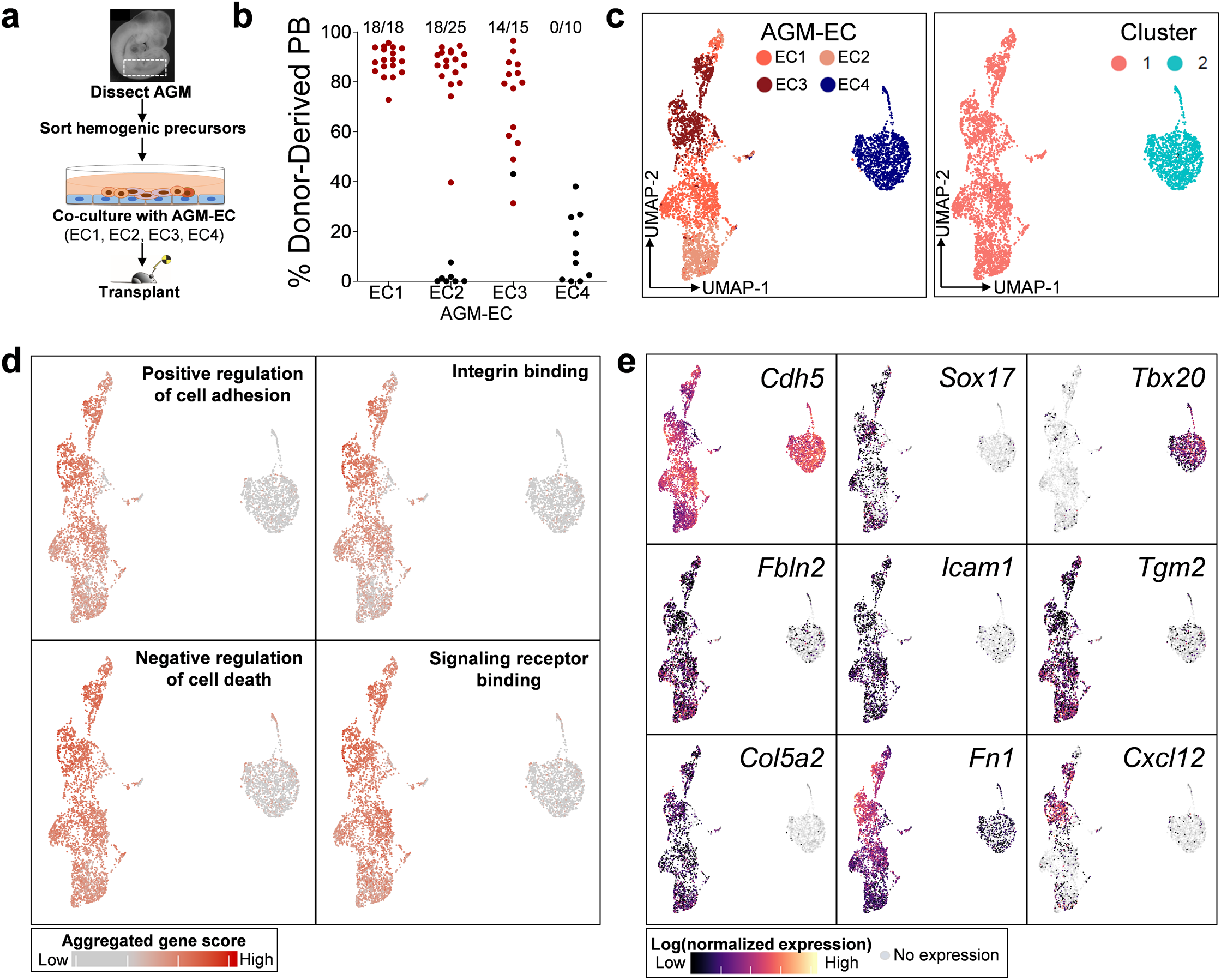
Single cell RNA-sequencing identifies the transcriptional signatures of AGM-EC that differentially support HSC generation in vitro. **a**. Methodology for comparison of AGM-EC in supporting generation of engrafting HSC from E9-10 AGM-derived VE-cadherin^+^ haemogenic precursors. **b**. Donor-derived peripheral blood (PB) engraftment in recipients transplanted with progeny of haemogenic precursors following co-culture on various independently generated AGM-EC (EC1-4). Numbers above indicate fraction of mice with multilineage engraftment, designated by data points in red. (PB engraftment shown at ≥24 weeks post-transplant, pooled from n≥3 independent experiments for each AGM-EC; see also Supplementary Table 1). **c**. UMAP and cluster analysis of single cell transcriptional profiles of each AGM-EC from (**b**). (See also Supplementary Fig. 1). **d**. Gene set scores representing top GO terms for transcripts differentially expressed by HSC-supportive verses non-supportive AGM-EC (biological processes and molecular functions ranked by p-value, see also Supplementary Table 2, Supplementary Table 3). **e**. Gene expression heatmap for pan-endothelial marker, VE-Cadherin (*Cdh5*), and selected transcripts differentially expressed between clusters. (See also Supplementary Table 2).

### VE-Cadherin^+^CD61^+^EPCR^+^ surface phenotype enriches for functional HSC precursors during the transition from HE to HSC in the AGM

Identifying signaling ligands from the transcriptomic analysis of AGM-EC that have functional relevance to HSC formation requires complementary knowledge of the cognate receptors and downstream pathways activated in haemogenic precursors during HSC specification and self-renewal. To capture this information, we next sought to characterize the single cell transcriptomes of haemogenic precursors during their transition from HE to HSC. However, the ability to isolate bona fide HSC precursors during early murine development is currently limited by the lack of unique markers which distinguish these precursors from more abundant non-haemogenic endothelial cells and haematopoietic progenitors lacking HSC potential^14, 19^. Using the AGM-EC co-culture system to screen for surface markers that enrich for HSC precursor activity, we previously demonstrated that HSC precursors from E9 to E11 are primarily restricted to the subset of VE-Cadherin^+^ cells expressing EPCR, a marker of HSCs throughout development^21, 26, 27, 31, 32^. We further determined that co-expression of EPCR with CD61, a surface marker shown to enrich for HE^33^, defines the minor subset of total AGM-derived VE-Cadherin^+^ cells with the most robust long-term HSC potential, as measured by serial transplantation following AGM-EC co-culture of sorted populations from as early as E9 (Fig. 2a-b, Supplementary Fig. 2). We previously reported a method utilizing single cell index sorting of embryonic haemogenic precursors with AGM-EC co-culture followed by functional analysis in transplantation assays to characterize the phenotypic and engraftment properties of clonal HSC precursors^26, 27^. Using this approach, we confirmed that clonal AGM-derived VE-Cadherin^+^EPCR^+^ cells with HSC potential co-express CD61, and furthermore demonstrate that cells with VE-Cadherin^+^CD61^+^EPCR^+^ phenotype (hereafter referred to as V^+^61^+^E^+^) encompass HSC precursors at different stages of development during their transition from phenotypic HE (CD41^-^CD45^-^) to pro-HSC/pre-HSC I (CD41^+^CD45^-^) and pre-HSC II/HSC (CD45^+^)^16, 17, 34^ (Fig. 2c-d). Collectively, these results demonstrate that the V^+^61^+^E^+^ phenotype enables substantial enrichment of rare HSC precursors across their developmental spectrum from HE.

**Figure 2.**
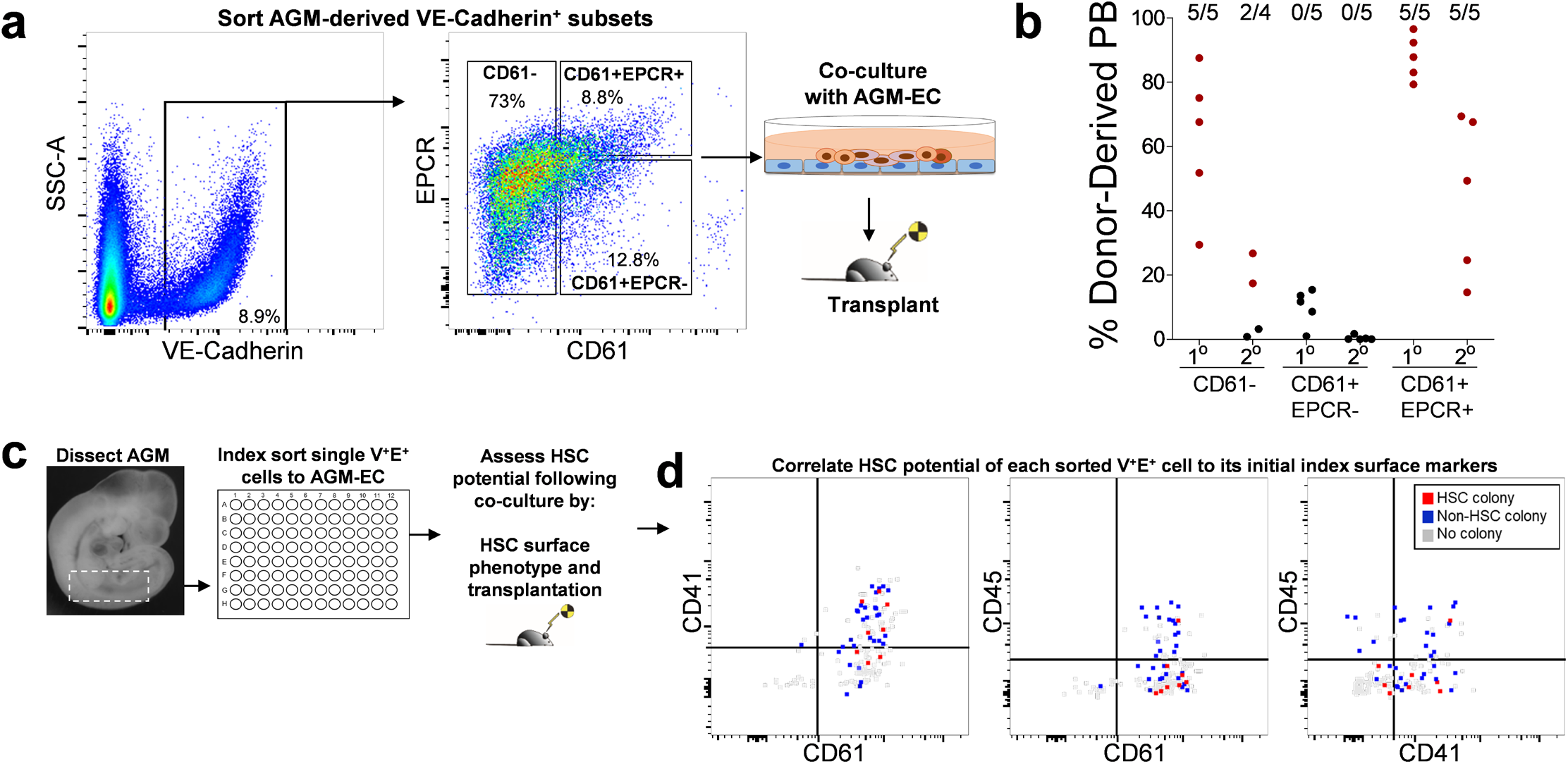
VE-Cadherin^+^CD61^+^EPCR^+^ (V^+^61^+^E^+^) surface phenotype enriches for functional HSC precursors during the transition from HE to HSC in the AGM. **a**. Sorting by FACS of E9 AGM/P-Sp (para-aortic splanchnopleura)-derived cells based on expression of CD61 and EPCR within the VE-Cadherin^+^ population (see also Supplementary Fig. 2). **b**. Each population in (**a**) was co-cultured on AGM-EC, transplanted to lethally irradiated adult mice, and analyzed for donor-derived peripheral blood (PB) engraftment in primary (20 weeks) and secondary (16 weeks) recipients. Numbers above indicate fraction of mice with multilineage engraftment in each group, designated by data points in red. (Similar results obtained in n=1 independent experiment. See also Supplementary Table 1). **c**. Methodology for index sorting of single VE-Cadherin^+^EPCR^+^ (V^+^E^+^) cells for co-culture on AGM-EC and subsequent analysis to assess HSC potential by flow cytometry and transplantation. **d**. Correlation of clonal HSC potential with surface expression of CD61, CD45 and CD41 on individual index sorted V^+^E^+^ cells. HSC colony (red) indicates formation of haematopoietic colony with HSC activity as detected by surface phenotype (VE-Cad^-/low^CD45^+^Gr1^-^F4/80^-^Sca1^hi^EPCR^hi^) and long-term (≥24 week) multilineage engraftment (see also Supplementary Table 1). Non-HSC colony (blue) indicates formation of haematopoietic colony lacking HSC phenotype and/or long-term engraftment. No colony (gray) indicates absence of detectable haematopoietic cells following AGM-EC co-culture.

### Single cell RNA-sequencing of AGM V^+^61^+^E^+^ cells identifies dynamic transcriptional signatures of HSC precursors during their transition from arterial-like HE

To study the transcriptional profiles of HSC precursors during their emergence from HE, we next isolated V^+^61^+^E^+^ cells by FACS from dissected AGM and performed scRNA-seq. Given HSC precursors are exceedingly rare prior to E10, we focused on AGM dissected between E10 and E11, during the peak emergence of HSC precursors from HE^14, 26^. We obtained scRNA-seq data from 3092 cells isolated from a total of 100 embryos pooled in two independent experiments at E10 (31-39 somite pairs) and E11 (40-45 somite pairs) (Supplementary Fig. 3a-e). Dimensionality reduction and unsupervised clustering of the single cell transcriptomes showed that most cells reside in a series of adjacent clusters (1, 2, and 5) differentially expressing endothelial markers (*Cdh5*, encoding VE-cadherin, and *Dll4*, encoding Notch ligand Delta-like-4, expressed in arterial EC) and haematopoietic fate-specifying transcription factors (*Runx1* and *Gfi1*), consistent with cells undergoing a transition from an endothelial to haematopoietic transcriptional program (Fig. 3a-b). Cell type classification based on marker genes identified that the major clusters (1, 2, and 5) are comprised of cells primarily expressing markers of arterial EC and haematopoietic fates (Fig. 3c, Supplementary Fig. 3f). Several smaller clusters segregated in transcriptional space express markers of differentiated myeloid and erythroid/megakaryocyte lineages (cluster 6), non-arterial EC (cluster 3), and somitic mesoderm (cluster 4), likely representing minor contaminating populations following FACS, which were excluded for subsequent analyses (Fig. 3c, Supplementary Fig. 3).

**Figure 3.**
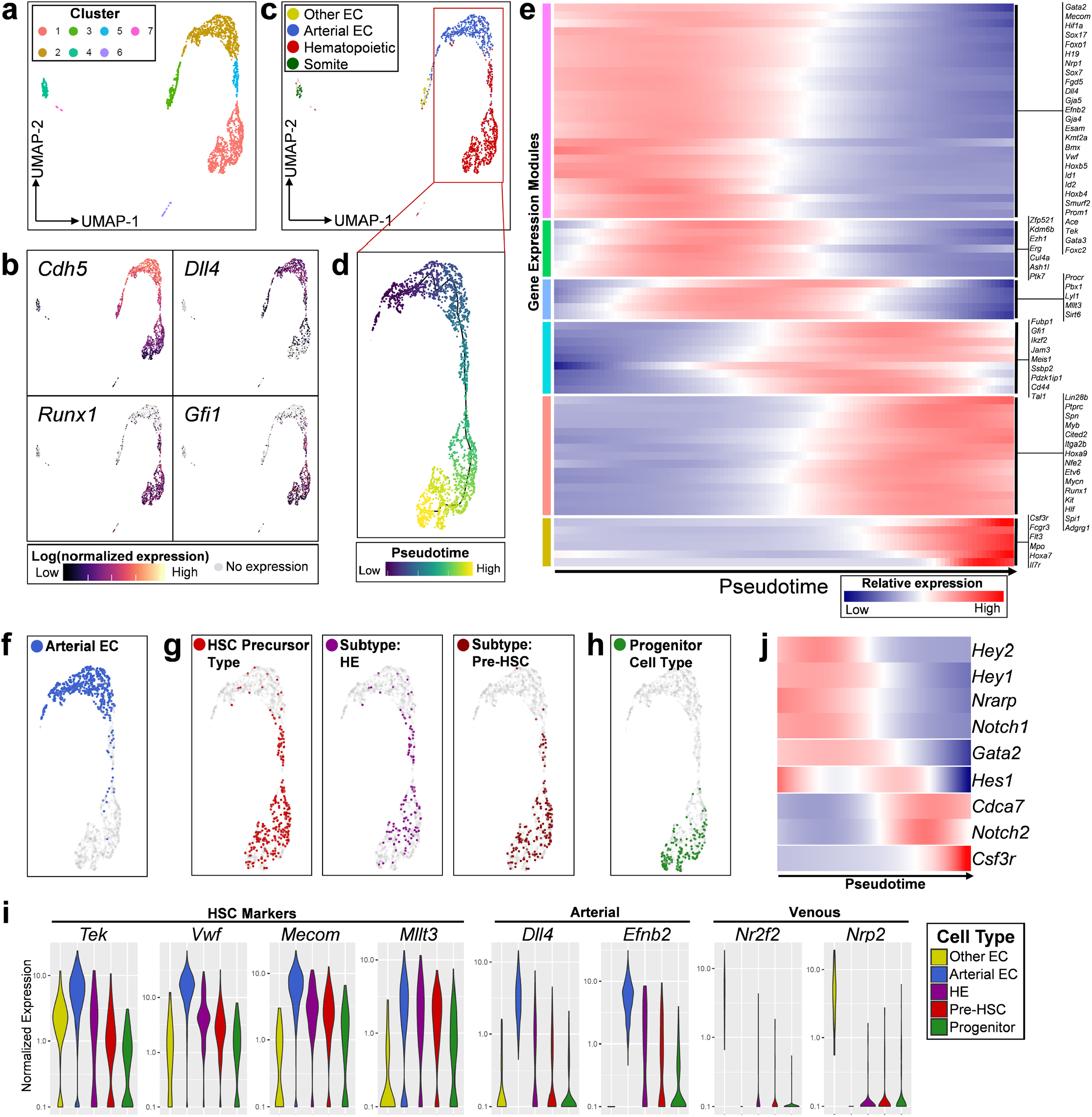
Single cell RNA-sequencing of AGM V^+^61^+^E^+^ cells identifies dynamic transcriptional signatures of HSC precursors during their transition from arterial-like HE in vivo. **a**. UMAP and cluster analysis of sorted E10 to E11 AGM-derived V^+^61^+^E^+^ cells. **b**. Expression heatmap of representative pan-endothelial (*Cdh5*/VE-Cadherin), arterial (*Dll4*), and haematopoietic-specific (*Runx1, Gfi1*) genes. **c**. Cell type classification of single cells based on marker genes (See also Supplementary Fig. 3f). **d**. Trajectory analysis using Monocle to order cells from adjacent clusters (1, 2, and 5) in pseudotime. **e**. Gene expression heatmap of selected differentially expressed genes over pseudotime, segregated by modules of gene expression (See also Supplementary Table 4). **f-h**. Cell type classification of single cells based on gene expression (See also Supplementary Fig. 3f): **f**. Arterial EC (*Cdh5, Dll4, Efnb2, Hey1*, not expressing *Runx1, Nr2f2, Nrp2*), **g**. HSC precursor type (*Runx1, Ctnnal1, Pdzk1ip1*, not expressing *Flt3, Il7r, Fcgr3, Csf3r*), and HSC precursor subtypes defined as HE (negative for *Itga2b*/CD41 and *Ptprc*/CD45) and pre-HSC (expressing *Itga2b* and/or *Ptprc*) **h**. Progenitor cell type (expressing *Runx1* plus one or more of *Flt3, Il7r, Fcgr3*, or *Csf3r*). **i**. Expression of HSC-marker and self-renewal genes (*Tek, Vwf, Mecom, Mllt3*), arterial EC genes (*Dll4, Efnb2*), and venous EC genes (*Nr2f2, Nrp2*) in individual cells defined by cell type classification from (f-h). **j**. Gene expression heatmap of Notch receptors and target genes in pseudotime. (See also Supplementary Fig. 3, Supplementary Table 4).

We then applied an unbiased machine learning algorithm using Monocle^35^ to generate a trajectory by pseudotemporal ordering of single cells (Fig. 3d) and to identify genes that vary significantly across pseudotime (Supplementary Table 4). Gene expression changes over pseudotime suggest this trajectory recapitulates the developmental emergence of pre-HSC/HSC (enriched in HSC-associated genes, egs. *Pdzk1ip1, Hlf, Etv6*) from an initial population expressing genes consistent with arterial EC/HE transcriptional programs (egs. *Dll4, Gja5, Nrp1, Sox17, Foxc2, Gata2*) (Fig. 3e). Notably, many genes associated with HSC-self renewal (egs. *Erg, Kdm6b, Mllt3, Mecom, Pbx1, Sirt6*)^36-41^ are most highly expressed in modules of gene expression in early pseudotime that overlap with arterial EC-specific programs and precede modules containing canonical haematopoietic-specifying transcription factors *Runx1* and *Gfi1*. Cells at the terminal portion of the pseudotemporal trajectory demonstrate changes in gene expression consistent with loss of HSC potential and acquisition of lymphoid/myeloid progenitor fates (egs. expression of *Flt3, IL7r, Csf3r, Fcgr3*) (Fig. 3e).

We next defined an “HSC precursor” cell type based on strict co-expression in single cells of haematopoietic-specific transcription factor *Runx1* together with *Ctnnal1* and *Pdzk1ip1* (HSC-marker genes that are not broadly expressed in endothelial cells) and absence of expression of progenitor-specific genes (*Flt3, IL7r, Csf3r*, and *Fcgr3*) (Fig. 3g)^42, 43^. Within this population of cells defined as “HSC precursors”, we confirmed the presence of subtypes transcriptionally resembling both “HE” (lacking expression of *Itga2b*/CD41 and *Ptprc*/CD45) and “pre-HSC” (expressing *Itga2b* and/or *Ptprc*), which are enriched in sequential portions of pseudotime consistent with their expected developmental progression (Fig. 3g, Supplementary Fig. 3f). We also defined a “progenitor” cell type based on co-expression of *Runx1* together with one or more progenitor-specific genes *Flt3, IL7r, Csf3r*, or *Fcgr3*; expectedly, cells of this type were largely confined to the terminal portion of pseudotime (Fig. 3h). Cells defined by “HSC precursor” type co-express other established HSC marker and self-renewal genes (egs. *Tek, Vwf, Mecom, Mllt3*) that were also highly expressed in “arterial EC” but lower in “progenitor” cell types (Fig. 3i). A subset of “HSC precursor” cell types also co-express genes specific to arterial fates (egs. *Dll4, Efnb2*) but largely lack expression of venous-specific genes (egs. *Nr2f2, Nrp2*) (Fig. 3i). Altogether, our single cell transcriptional data, comprising a highly enriched population of functionally validated HSC precursors, combined with further in silico clustering and trajectory analysis, successfully captures the developmental emergence of HSCs from HE with an arterial endothelial transcriptional program.

By distinguishing the transcriptional profiles of the rare transitional population of emerging HSC precursors from more differentiated haematopoietic progenitors, this analysis enables us to identify receptors and downstream signal pathways specifically expressed during pre-HSC/HSC development from HE. For example, to study the dynamics of Notch signaling during this process, we examined relative expression of genes activated by the Notch pathway over pseudotime (Fig. 3j). During the maturation of HE to pre-HSC/HSC, concomitant with decreasing *Notch1* and increasing *Notch2* expression, expression of Notch target genes *Hey1/Hey2*, required for arterial EC fates^44^, are downregulated, and Notch target gene *Cdca7*, which regulates HSC emergence^45^, is increased. Notch targets *Gata2* and *Hes1*, also essential for HSC development^46-48^, are sequentially downregulated in the terminal portion of pseudotime corresponding with increasing expression of genes specific to differentiation to progenitors (*Csf3r*). This pattern of Notch pathway gene expression changes is consistent with recent studies suggesting that, although Notch1 is required early for definitive haematopoietic fate, emergence of HSCs from HE correlates with decreased *Notch1* expression, lower overall Notch signal strength, and a requirement for *Hes1* in limiting Notch-dependent *Gata2* expression^6, 48-51^. These results demonstrate the utility of our single cell transcriptomic analysis to elucidate the developmental dynamics of signaling pathways during the HE to pre-HSC/HSC transition.

### Single cell RNA-sequencing identifies the transcriptional signatures of self-renewing HSCs emerging from haemogenic precursors in vitro

We previously showed that a single AGM-derived haemogenic precursor could give rise to more than one hundred long-term engrafting HSCs following AGM-EC co-culture, suggesting that the AGM-EC niche provides signals sufficient to support both the maturation and self-renewal of HSCs^26^. Therefore, we hypothesized that single cell transcriptomic analysis of HSCs generated during AGM-EC co-culture would identify receptors and downstream signal pathways essential to HSC formation and self-renewal in vitro, complementing our analysis of primary haemogenic precursors undergoing the initial transition from HE to pre-HSC/HSC in vivo. Thus, we next sought to acquire the transcriptional profiles of HSCs emerging from embryonic haemogenic precursors in the AGM-EC vascular niche by performing scRNA-seq on the progeny of individual FACS-isolated AGM-derived V^+^61^+^E^+^ cells during colony formation in AGM-EC co-culture. Cells were harvested at day 4 of co-culture, and 50% of the cells were analyzed for surface phenotype we previously showed to correlate with long-term engrafting HSC activity (VE-Cadherin^-/low^CD45^+^Gr1^-^F4/80^-^Sca1^hi^EPCR^hi^)^26, 27^. scRNA-seq was performed from the remaining cells from two colony types, one with a relatively homogeneous population of phenotypic HSCs (Fig. 4a) and another consisting of a mixed population of phenotypic HSCs and cells with decreasing expression of Sca1 and EPCR consistent with differentiation to haematopoietic progenitor cells (HPC) (Fig. 4b). Dimensionality reduction and clustering analysis identified distinct clusters consistent with haematopoietic progeny (expressing *Ptprc/*CD45) for each colony type (Fig. 4c-d, Supplementary Fig. 4). Consistent with their HSC phenotype, the cells from the haematopoietic cluster in the first colony type uniformly expressed established HSC marker genes (such as *Pdzk1ip1* and *Vwf*) and largely lacked expression of genes associated with differentiation of HSC to HPC (*Itgal/*CD11a, *Cd48*)^52, 53^ (Fig. 4e). Haematopoietic cells in the second colony type demonstrated a similar, minor population of HSCs based on their transcriptional signatures (“HSC Type”, co-expressing *Runx1, Pdzk1ip1* and *Vwf*, negative for *Cd48* and *Itgal*) and a larger population of emerging haematopoietic progenitor cells (“HPC Type”, expressing *Cd48* and/or *Itgal*) (Fig. 4f). Pseudotemporal ordering recapitulated the differentiation of HPC from HSC, revealing gene expression changes during this process, including increased expression over pseudotime of genes involved in early myeloid and lymphoid-specific fates, egs *Cebpa* and *Ebf1* (Fig. 4g, Supplementary Table 5). Consistent with our previous studies showing that both emergence of HSCs from HE and expansion of HSCs in the vascular niche are Notch-dependent^25, 54^, analysis of scRNA-seq data from HSCs generated during co-culture with AGM-EC demonstrates that Notch target genes required for HSCs (*Hes1, Gata2*) are enriched in cells in early pseudotime expressing HSC-specific programs, including genes involved in HSC self-renewal, such as *Cdkn1c* (p57), *Mecom, Pbx1*, and *Mllt3*, with downregulation concomitant with differentiation to HPC (Fig. 4g). Collectively, this scRNA-seq data captures the transcriptomes of HSCs during their emergence from clonal V^+^61^+^E^+^ precursors and reveals the dynamics of HSC self-renewal and early HSC differentiation in the AGM-EC niche.

**Figure 4.**
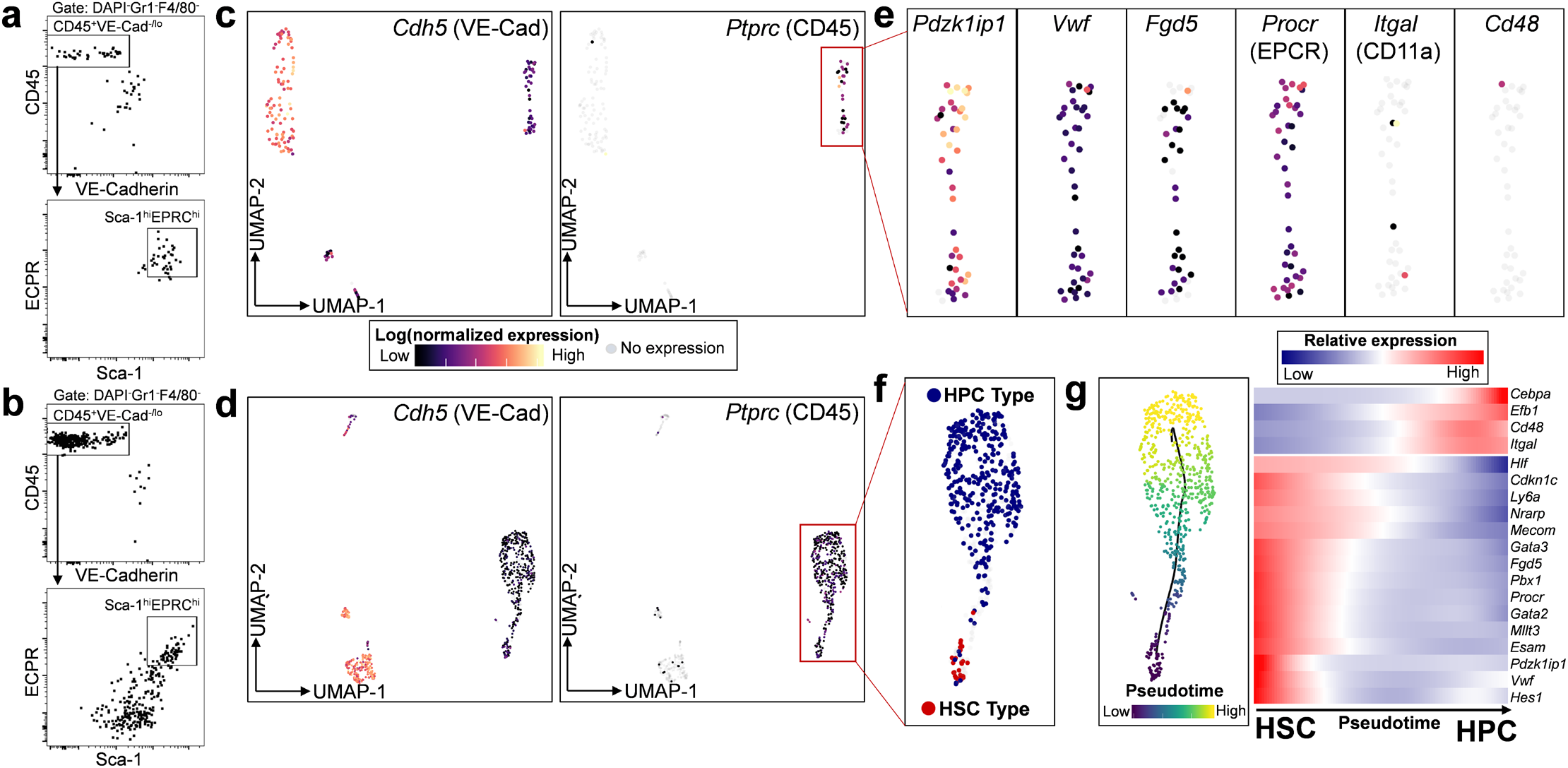
Single cell RNA-sequencing identifies the transcriptional signatures of self-renewing HSCs emerging from clonal V^+^61^+^E^+^ haemogenic precursors in the AGM-EC niche in vitro. **a-b**. FACS analysis of the progeny of a single V^+^61^+^E^+^ haemogenic precursor following co-culture on AGM-EC, with (**a**) HSC phenotype (VE-Cad^low/-^CD45^+^Gr1^-^F4/80^-^Sca1^hi^EPCR^hi^) and (**b**) mixed HSC and haematopoietic progenitor cell (HPC) phenotype. **c-d**. UMAP projection of single cell transcriptomes obtained from cells from a-b showing expression of endothelial marker *Cdh5* (VE-cadherin) and pan-haematopoietic marker *Ptprc* (CD45). **e**. Expression of HSC-marker genes (*Pdzk1ip1, Vwf, Procr, Fgd5*) and HPC-associated genes (*Itgal, Cd48*) in haematopoietic cluster (expressing *Ptprc*). **f**. Cell type classification of haematopoietic cells based on gene expression: HSC Type (expressing *Runx1, Pdzk1ip1*, and *Vwf*, not expressing *Cd48* or *Itgal*) and HPC type (expressing *Cd48* and/or *Itgal*). **g**. Trajectory analysis using Monocle to order cells from the haematopoietic cluster in pseudotime, and gene expression heatmap over pseudotime of selected genes, including Notch pathway targets, differentially expressed during HSC differentiation to HPC. (See also Supplementary Fig. 4, Supplementary Table 5).

### Complementary analysis of the transcriptional profiles of AGM-EC and developing HSCs identifies ligand-receptor interactions supporting HSC development

For the AGM-EC niche to support the generation of functional HSCs from HE, it must express a repertoire of signaling ligands necessary to promote both the maturation and self-renewal of HSC. In parallel, developing HSCs must express the cognate receptors required to receive these essential niche-derived signals. Thus, we hypothesized that analysis of the complementary transcriptomes of AGM niche EC and HE/pre-HSCs transitioning to self-renewing HSCs from our in vivo and in vitro scRNA-seq data could be used to identify the ligand-receptor interactions necessary to support HSC development. To test this, we applied a comprehensive database consisting of curated pairs of ligands and receptors^55^ to our scRNA-seq data to infer potential functionally relevant interactions. We first identified ligands expressed in the HSC-supportive AGM-EC (from Fig. 1), as well as primary AGM-derived arterial EC (“Arterial EC” Type, from Fig. 3c), which share common phenotypic characteristics and overlapping gene expression profiles (Supplementary Fig. 5). We then identified receptors expressed in the subset of AGM-derived V^+^61^+^E^+^ cells transcriptionally defined as “HSC precursor” type (from Fig. 3g), representing cells encompassing the transition from HE to pre-HSC/HSC. These data sets were then combined to determine the global set of ligand-receptor interactions between AGM niche EC and HE/pre-HSC (Fig. 5, Supplementary Table 6). We also identified receptors expressed in cells defined by “HSC Type” emerging during AGM-EC co-culture of AGM-derived V^+^61^+^E^+^ cells (from Fig. 4) and combined this data with AGM niche EC ligands to determine the global set of ligand-receptor interactions potentially regulating HSC maturation and self-renewal (Fig. 5, Supplementary Table 6). We used an unbiased identification of ligands and receptors based only on expression thresholds in each cell type, so as to provide a comprehensive list of potential interactions (see Methods). However, ligand-receptor interactions that involved ligands enriched in HSC-supportive verses non-supportive AGM-EC are highlighted (from Supplementary Table 2, indicated by yellow arrows in Fig. 5), as we hypothesize that these interactions may account for functional differences in AGM-EC niche support of HSCs. This integrated analysis identified receptor-ligand interactions modulating signaling pathways with established functions in HSC development and self-renewal, such as Notch and Wnt^5, 56-58^, as well as potential novel interactions involving integrins, cell adhesion molecules, cytokines/chemokines, growth factors, and guidance signal pathways (Fig. 5i-viii).

**Figure 5.**
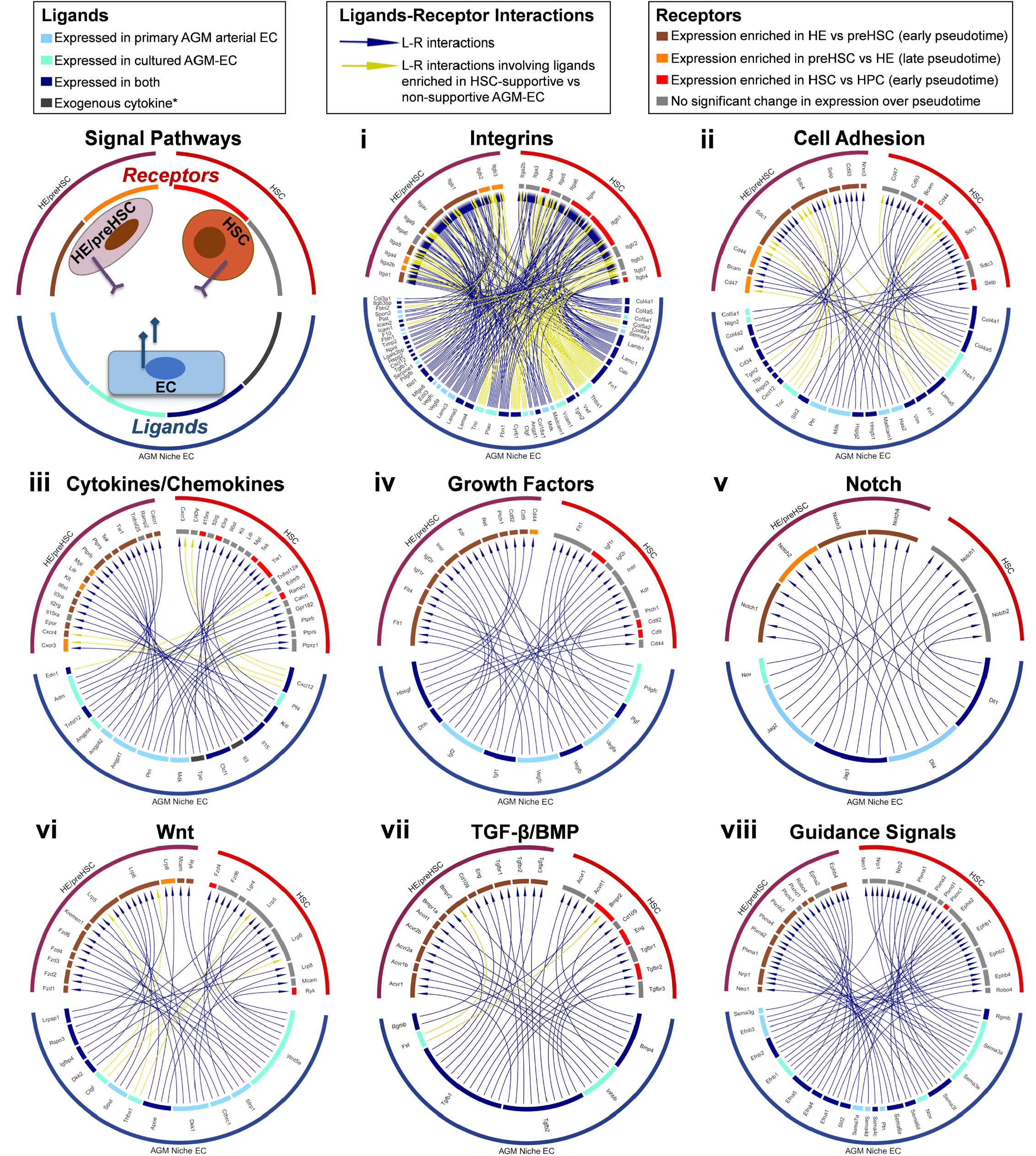
Signaling interactions identified by ligand-receptor analysis. Selected signal pathways regulated by ligand-receptor interactions between AGM niche EC and developing HSCs based on complementary scRNA-seq analysis. HE/PreHSC: Indicates receptors expressed in primary AGM V^+^61^+^E^+^ cells classified as “HE” or “pre-HSC” types (see Fig. 3g). HSC: Indicates receptors expressed in cells generated following AGM-EC co-culture of V^+^61^+^E^+^ cells classified as “HSC” type (see Fig. 4). Receptors were also classified as to whether they were expression more highly in early (enriched in HE) or late (enriched in pre-HSC) pseudotime for primary AGM V^+^61^+^E^+^ cells, or expressed more highly in early pseudotime (enriched in HSCs vs HPCs) in cells generated following AGM-EC co-culture. Ligands were classified by whether they were expressed in HSC-supportive AGM-EC (cultured AGM-EC), primary AGM-derived AGM V^+^61^+^E^+^ cells classified as “arterial EC” (primary AGM arterial EC), or both. Signals mediated by ligands differentially expressed in HSC-supportive vs non-supportive AGM-EC are indicated by yellow arrows (see Supplementary Table 2). *indicates exogenous cytokines in AGM-EC culture conditions. (See also Supplementary Table 6 for complete list of identified ligands, receptors, and ligand-receptor interactions).

### Functional analysis of ligand-receptor interactions enables the engineering of an in vitro niche supporting HSC development from haemogenic precursors

Building from the ligand-receptor interactions identified by our single cell transcriptomic analysis, we next sought to engineer an artificial stroma-independent niche sufficient to recapitulate the function of AGM-EC in supporting the generation of long-term engrafting HSCs from embryonic HSC precursors. We previously identified a set of conditions incorporating immobilized Notch ligand Delta1 (Dll1-Fc) that was sufficient to support quantitative expansion of HSCs isolated from E11 AGM, at which time in development engrafting HSCs are already detectable at low frequency by direct transplantation into adult recipients^25^. However, these conditions were unable to support the generation of engrafting HSCs from haemogenic precursors isolated prior to E11. We adapted these conditions by incorporating serum-free media to further control for exogenous factors. Based on cytokine receptor expression profiles in the scRNA-seq data in AGM-derived HE/pre-HSC and in vitro generated HSCs (Fig. 5iii, Supplementary Table 6), we retained recombinant stem cell factor, interleukin-3, and thrombopoietin. Based on our single cell transcriptomic analysis and reported studies suggesting dynamic roles for Notch1 and Notch2 receptors in activating downstream Notch targets expressed during pre-HSC/HSC formation from HE and subsequent HSC maturation and self-renewal^6, 25, 49-51, 59, 60^ (Fig. 3j, Fig. 4g, Fig. 5v), we also incorporated immobilized anti-Notch1 and anti-Notch2 monoclonal antibodies in place of immobilized Dll1-Fc. These antibodies specifically bind Notch1 and Notch2 receptors, respectively^25^, enabling targeted activation of Notch1 and/or Notch2 receptors independent of Fringe-mediated modifications that could alter their binding and activation by ligands such as Dll1^61^. Based on the differential expression of *Fn1* (fibronectin) in HSC-supportive vs non-supportive AGM-EC, and its interaction with integrins including VLA-4 (*Itga4*/*Itgb1*) and VLA-5 (*Itga5*/*Itgb1*) identified in our receptor-ligand analysis (Fig. 1, Fig. 5i), we also utilized immobilized recombinant fibronectin fragment (FN-CH-296), which specifically binds cell-surface VLA-4/VLA-5 in vitro.

We then tested whether components of this engineered niche including immobilized FN-CH-296, cytokines (IL3, SCF, and TPO), and small molecule inhibitor of TGF-β receptor (LY364947), and either immobilized Dll1-Fc, anti-N1 and anti-N2 antibodies (aN1/N2 Ab), or IgG (control), could support in vitro generation of HSCs. Long-term engrafting HSCs were detected following culture of E11 AGM-derived V^+^61^+^E^+^ cells in engineered conditions with either immobilized Dll1-Fc or combined aN1/N2 antibodies, but not control IgG (Fig. 6a-b, Supplementary Fig. 6a-c). However, these conditions were unable to support generation of engrafting HSCs from V^+^61^+^E^+^ cells isolated from AGM prior to E11 (Fig. 6c and data not shown), suggesting additional signals are required to promote the maturation of HSCs from haemogenic precursors at earlier stages.

**Figure 6.**
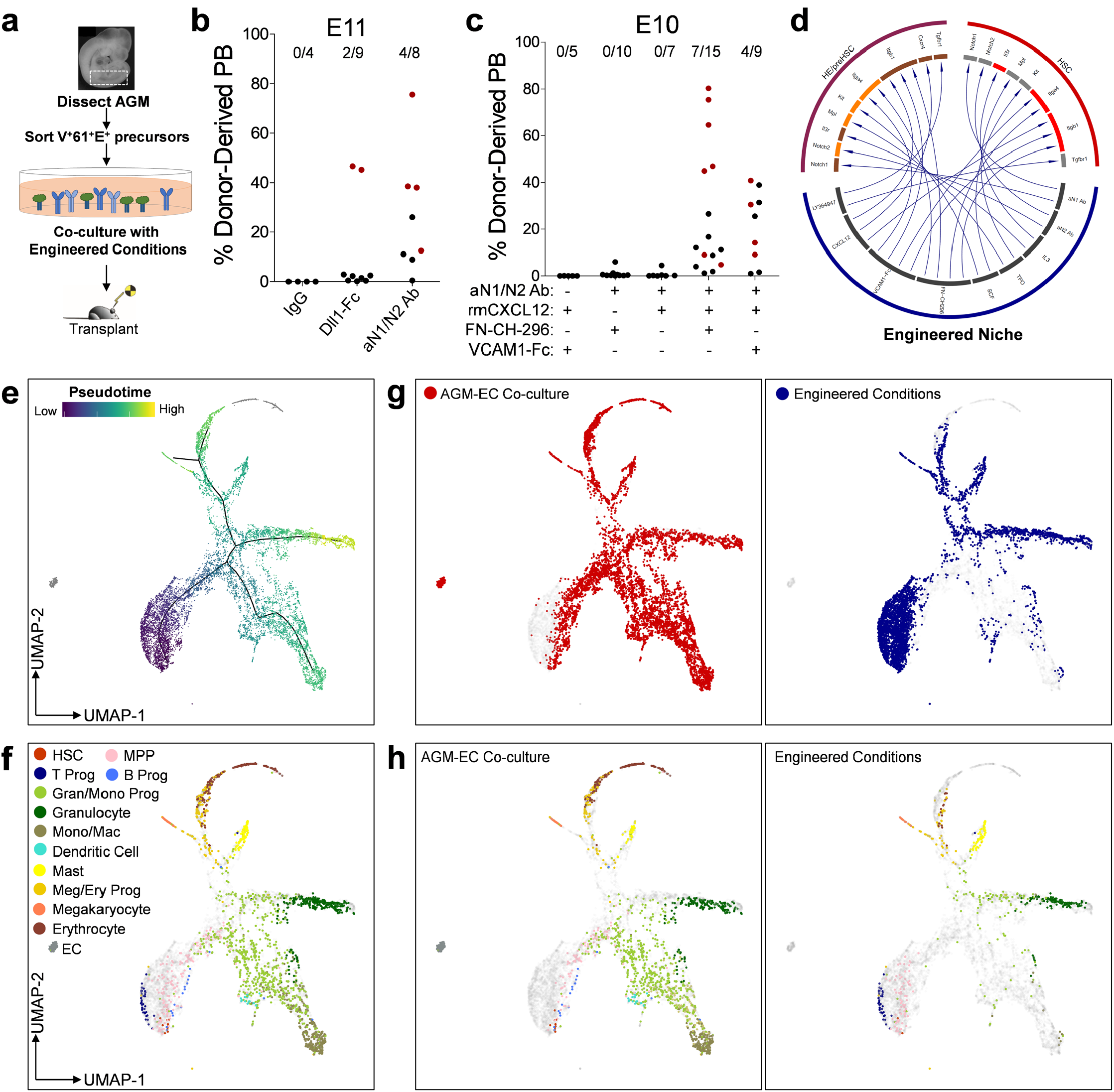
Functional validation of ligand-receptor interactions enables the engineering of an in vitro niche supporting HSC development. **a**. Methodology evaluating generation of engrafting HSC from AGM-derived V^+^61^+^E^+^ haemogenic precursors in engineered conditions. **b**. Donor-derived peripheral blood engraftment (≥24 weeks) in recipients transplanted with progeny of E11 AGM-derived V^+^61^+^E^+^ haemogenic precursor following co-culture in engineered conditions with immobilized Dll1-Fc, anti-Notch1/Notch2 (aN1/N2 Ab), or control (IgG). Data points in red indicate mice with multilineage engraftment (results pooled from n=2 independent experiments; see also Supplementary Fig. 6, Supplementary Table 1). **c**. Donor-derived peripheral blood engraftment (≥24 weeks) in recipients transplanted with progeny of E10 AGM-derived V^+^61^+^E^+^ haemogenic precursor haemogenic precursor following co-culture in engineered conditions as indicated (results pooled from n=5 independent experiments). **d**. Ligand-receptor interactions recapitulated in the engineered niche. **e**. UMAP and pseudotemporal ordering of single cell transcriptomes of the progeny of E10 V^+^61^+^E^+^ haemogenic precursors following 6 days of culture on AGM-EC or engineered conditions. **f**. Cell type classification based on marker genes. G-H. Subsets of single cell transcriptomes derived from cells cultured on AGM-EC or engineered conditions. (See Supplementary Fig. 6).

To identify signals required for the initial steps in HSC maturation from early HSC precursors, we prioritized for further analysis ligand-receptor interactions involving ligands that were expressed in primary AGM arterial EC and also significantly enriched in HSC-supportive versus non-supportive AGM-EC (Fig. 5, ligands indicated in navy, interactions indicated by yellow arrows), and to cognate receptors in HSC precursors whose expression is enriched in early pseudotime during the HE to pre-HSC/HSC transition (Fig. 5, receptors indicated in dark red). This identified genes encoding cell adhesion and extracellular matrix proteins that interact with integrin receptors, including *Fn1* (fibronectin) and *Tgm2* (tissue transglutaminase, which can enhance integrin β1 interaction with fibronectin)^62^, and genes encoding secreted factors *Cyr61*, an integrin-interacting protein, and *Cxcl12*, a chemokine that interacts with the receptor encoded by *Cxcr4* expressed in HE based on our scRNA-seq data (Fig. 5iii, Supplementary Table 6). Amongst exogenous ligands tested, we found that addition of recombinant CXCL12 reproducibly enabled the generation of multilineage long-term engrafting HSCs from E10 AGM-derived V^+^61^+^E^+^ cells following in vitro culture (Fig 6c and data not shown). Generation of HSCs in the presence of CXCL12 at this stage was also dependent on Notch activation by immobilized aN1/N2 Ab and the presence of a ligand for integrin VLA-4, either immobilized fibronectin fragment (FN-CH-296) or VCAM1-Fc chimera (Fig. 6c). In some experiments, these engineered conditions were sufficient to support generation of engrafting HSCs from V^+^61^+^E^+^ cells derived from embryos as early as E9, though with lower efficiency than at later stages (Supplementary Table 1).

In addition to supporting the generation and self-renewal of HSCs from haemogenic precursors, we previously showed that co-culture in the AGM-EC niche supports multilineage haematopoietic differentiation in vitro^25^. To compare the emergence of HSCs and downstream haematopoietic lineages in the engineered niche with AGM-EC co-culture, we performed scRNA-seq on the haematopoietic progeny generated following extended culture of AGM-derived V^+^61^+^E^+^ cells in each condition. Haematopoietic progeny following culture in the engineered niche included rare cells with transcriptional signatures of HSC and multipotent progenitors, as expected, as well as cells differentiating to myeloid, erythroid/megakaryocytic, and lymphoid lineages (Fig. 6e-h, Supplementary Fig. 6). Relative skewing of contribution toward early T lymphoid fate and away from B lymphoid, monocyte/macrophage, and late erythroid fates in engineered conditions suggests relatively increased activation of Notch pathway and/or deficiency of yet undefined signals presented by AGM-EC stroma. Collectively, these results demonstrate that a stroma-independent engineered niche is sufficient to partially recapitulate the vascular niche supporting HSC development from embryonic haemogenic precursors, and that further optimization through incorporation of additional niche-derived signals and temporal modulation of key signal pathways such as Notch during culture could further enhance the balance of HSC generation and self-renewal versus lineage differentiation.

## DISCUSSION

In this study, we provide several advances towards deciphering the complex signaling interactions required to support the specification and self-renewal of HSCs during embryonic development. First, using an ex vivo vascular niche modeling the embryonic AGM, where the first HSCs originate from HE, we identified a combination of phenotypic markers (V^+^61^+^E^+^) that enable substantial enrichment of functionally-validated HSC precursors encompassing the transition from HE to pre-HSC/HSCs. Second, using scRNA-seq of the primary AGM V^+^61^+^E^+^ population and the progeny of these cells during their maturation to HSCs and self-renewal in the AGM-EC niche, we generated a comprehensive transcriptional map of HSC development from HE. Third, we utilized this data to identify receptors and signal pathways expressed during HSC specification and self-renewal and, combined with complementary transcriptional analysis of the HSC-supportive AGM EC niche to identify cognate ligands, produced an atlas of potential ligand-receptor interactions regulating HSC development in the vascular niche. Finally, we applied this knowledge to rationally design a stromal cell-independent engineered niche sufficient to generate engrafting HSC from embryonic haemogenic precursors in vitro.

Our approach to single cell transcriptomic analysis of HSC development in the AGM builds on recently reported studies^21-23^, providing a comprehensive transcriptional map of emergence of HSCs from an earlier step in their developmental progression and adding critical information about interactions with niche ECs supporting the process of HSC specification and self-renewal. Because we functionally validated HSC precursor activity within the V^+^61^+^E^+^ population independent of CD41 and CD45 expression, we were able to capture the transcriptional profiles of V^+^61^+^E^+^CD41^-^CD45^-^ HE that precede the emergence of V^+^61^+^E^+^CD41^+^CD45^+/-^ pre-HSC I/II. Our analysis, combining functional, phenotypic, and transcriptional studies at the single cell level, thus provides strong evidence that HSCs emerge from HE with an arterial endothelial transcriptional program, supporting recent studies from our group and others that suggest multilineage definitive haematopoiesis and HSC potential resides within HE/pre-HSCs with arterial endothelial phenotypes^8, 10, 26^. Intriguingly, during the transition from arterial-like EC/HE to pre-HSC/HSC represented by pseudotemporal ordering of single cell transcriptomes, the expression of multiple genes required for HSC maintenance and self-renewal (egs. *Mllt3, Erg, Pbx1, Mecom, Sirt6, Cul4a, Ptk7, Kdm6b, Zfp521, Ezh1*)^36-41, 63, 64^ precedes the upregulation of transcription factors *Runx1* and *Gfi1*, which are required to specify haematopoietic fate (Fig. 3e), suggesting that self-renewal programs linked to arterial endothelial/HE states may be uniquely required prior to hematopoietic-fate specification for the generation of HSCs. Our studies thus suggest that recapitulating the conditions necessary to promote these arterial endothelial/HE programs in vitro will be a critical initial step to establishing methods for generating HSCs from pluripotent stem cells. Further supporting this concept, an elegant study of HSC development combining scRNA-seq with single cell ATAC-seq, reported during the preparation of this manuscript, identified a population of putative precursors to HE characterized by high expression of arterial genes such as *Sox17* and *Hey2*, in which multiple hematopoietic stem and progenitor cell-associated transcriptional programs appear to converge to regulate *Runx1* expression^*65*^.

Our study also leveraged an AGM-derived EC niche model, which we previously showed could support the generation and expansion of long-term engrafting HSCs from embryonic precursors^25-27^, to identify niche-derived ligands essential to HSC specification and self-renewal. We were able to identify signaling ligands with functional relevance based on their differential expression in AGM-EC that varied in their ability to support HSC generation from haemogenic precursors prior to E11. We hypothesize that heterogeneity in the supportive properties of AGM-EC stroma may result from founder effects due to the small population of AGM-derived EC that initially survives following FACS and cell culture to establish each line. Previous studies suggest that mesodermal origin and dorso-ventral positioning contribute to heterogeneity in the HSC-inductive properties of EC of the AGM aorta^66-68^. Interestingly, the non-supportive AGM-EC uniquely expressed *Tbx20*, a transcription factor previously shown to be expressed in endothelial cells localized to the dorsal portion of the embryonic aorta which lacks HSC activity (Fig. 1d)^30, 66^. In contrast, HSC-supportive AGM-EC shared expression of *Sox17*, a transcription factor required for arterial EC/HE fates, with primary arterial EC and HE (Fig. 1, Supplementary Fig. 5)^7, 29^. We hypothesize that V^+^61^+^E^+^ cells include non-haemogenic arterial ECs that deliver HSC-supportive signals to nearby V^+^61^+^E^+^ HE in the AGM. Supporting this hypothesis, primary AGM-derived V^+^61^+^E^+^ cells identified transcriptionally as arterial EC (Fig. 3c, f) exhibit a gene expression profile overlapping with that of HSC-supportive AGM-EC, including expression of essential niche ligands functionally validated in our engineered niche, such as *Cxcl12* and *Fn1*. Future studies will further determine how differential HSC-supportive properties of AGM-EC may relate to heterogeneity of the primary niche EC of the aorta from which they are derived.

By prioritizing ligand-receptor interactions that were specifically identified in HSC-supportive AGM-EC, our analysis highlighted the key role of integrins. While integrin interactions are known to mediate aspects of HSC homing, retention, and maintenance in fetal liver and adult bone marrow niches, a study in zebrafish also demonstrated an important role for fibronectin-integrin β1 interactions in promoting Notch-mediated HSC fate in the embryonic aorta^69^. Consistent with this study, we determined that the combination of Notch activation, mediated by immobilized Notch1 and Notch2-receptor specific antibodies, and a VLA-4 integrin (*Itga4*/*Itgb1*) ligand, either VCAM1-Fc chimera or Fibronectin fragment, was necessary to support HSC generation from embryonic AGM-derived haemogenic precursors in our engineered niche. Moreover, we identified a specific requirement for the chemokine CXCL12 in the engineered niche to enable the generation of engrafting HSCs from haemogenic precursors isolated prior to E11, the earliest developmental timepoint in which HSC activity is consistently detected by direct transplantation. This suggests an important role for the chemokine CXCL12 in promoting HSC fate from less mature haemogenic precursors. Consistent with this finding, a study in zebrafish demonstrated that a subset of aortic endothelial cells derived from somitic mesoderm supported HSC fate from aortic HE via production of CXCL12^68^. Expression of the canonical receptor for CXCL12, CXCR4, is detected in arterial EC and HE by our single cell transcriptional analysis, though its expression is downregulated during the transition from HE to pre-HSC/HSC. VLA-4 integrin has also been shown to be activated by CXCL12 by direct binding, potentiating its interactions with ligand VCAM1, suggesting an additional mechanism for CXCL12 enhancement of HSC generation in our engineered niche^70^. Futures studies will be required to determine the precise mechanism by which CXCL12 interacts with its receptor(s) to promote stage-specific aspects of HSC generation.

Altogether, our studies identified molecular interactions predicted by single cell transcriptomics to establish unique engineered conditions sufficient to support generation of engrafting HSCs from embryonic haemogenic precursors. Building on this genome-wide map of the ‘interactome’ regulating HSC development in the AGM vascular niche, future studies will be necessary to identify additional factors required to optimize HSC generation from earlier stages of development. Based on dynamic changes in expression of genes involved in key signal pathways, such as Notch, Wnt, and TGF-β, we hypothesize that stage-specific modulation of these pathways during culture in the engineered niche will enable further optimization of HSC specification and self-renewal. Furthermore, we hypothesize that signals derived from non-endothelial AGM niche cells, such as surrounding mesenchymal populations, may be critical to further enhance HSC generation. Altogether, our studies represent a significant step toward precisely defining the niche factors required for HSC generation, which will be critical for advancing therapeutic applications such as for disease modelling and cellular therapies from human pluripotent stem cells.

## METHODS

### Reagents

Source and catalog number for key reagents and antibodies are listed in Supplementary Table 7.

### Mice

Wild type C57Bl6/J7 (CD45.2) and congenic C57BL/6.SJL-Ly5.1-Pep3b (CD45.1) mice were bred at the Fred Hutchinson Cancer Research Center. C57Bl6/J7 CD45.2 mice were used for timed matings. All animal studies were conducted in accordance with the NIH guidelines for humane treatment of animals and were approved by the Institutional Animal Care and Use Committee at the Fred Hutchinson Cancer Research Center.

### Embryo dissections and cell sorting

Embryo AGM tissues (or P-Sp, para-aortic splanchnopleure - precursor region to the AGM prior to E10) were dissected from embryos harvested from pregnant C57Bl6/J7 (CD45.2) female mice as previously described^25^. Embryo age was precisely timed by counting somite pairs (sp): E9 (21-29sp), E10 (30-39sp), and E11 (40-47sp). Dissected AGM/P-Sp tissues were treated with 0.25% collagenase for 25 minutes at 37°C, pipetted to single cell suspension, and washed with PBS containing 10% FBS. Cells were incubated with anti-mouse CD16/CD32 (FcRII block) and stained with monoclonal antibodies as indicated. A detailed list of all antibodies used is shown in the Supplementary Table. For most experiments, to isolate haemogenic endothelial populations, a combination of anti-mouse VE-Cadherin-PECy7, EPCR/CD201-PerCP-eFluor710, and CD61-APC were used, with or without additional anti-mouse antibodies for CD45 (PE or FITC) and CD41 (PE or AF488), as indicated in results section. DAPI staining was used to gate out dead cells. All reagents for cell staining were diluted in PBS with 10% FBS and staining was carried out on ice or at 4°C. Cells were sorted on a BD FACSAria II equipped with BD FACSDiva Software with index sorting capability. For index-sorted single cells, sorting was performed in single cell mode with maximum purity mask settings to minimize contaminating cells.

### AGM-derived Akt-EC (AGM-EC) generation

AGM-EC were derived as previously described^25^. Briefly, AGM dissected from pooled E10–E11 embryos were sorted by FACS for endothelial cells as VE-cadherin^+^CD45^-^CD41^-^. Sorted cells were cultured on 48-well tissue culture plates (density >20,000 cells per well) pre-treated with 5 μg/ml RetroNectin (r-fibronectin CH-296), in EC media (consisting of IMDM with 20% FBS, Penicillin/streptomycin, L-glutamine, heparin 0.1 mg/ml, and endothelial mitogen 100 μg/ml). For initial culture, EC media was supplemented with VEGF (50 ng/ml), CHIR009921 (5 μM), and SB431542 (10 μM). Following 1–2 days of culture, surviving cells (which form scattered, small tightly adherent colonies) were infected by lentivirus with pCCL-PGK myrAkt construct, as previously described^71^. When confluent, cells were serially split to 24-well, 6-well, and T75 tissue culture plates (pre-treated with 0.1% gelatin) in EC media without added VEGF, CHIR09921, or SB431542. The first passage to T75 flask was considered passage 0, with subsequent passages every 3–4 days plated at approximately 5×10^5^ cells per T75 flask. All AGM-EC derivations were tested by plating cells at 4×10^4^ cells/24 wells in X-vivo 20 media to ensure that a confluent layer of viable ECs was maintained for at least 7 days in serum-free conditions. Endothelial identity/purity for each independently derived AGM-EC was confirmed by FACS for surface expression of VE-Cadherin. Multiple, independently derived AGM-EC were frozen at low passage at 1×10^6^ cells/vial for subsequent experiments. For co-culture experiments to compare HSC-supportive capacity, AGM-EC from multiple derivations were thawed in parallel from low passage aliquots onto gelatin-treated T75 flasks in EC media above, passaged every 3-4 days when confluent, and used at similar passage number for serum-free co-culture assays with freshly sorted haemogenic precursors, as described below. For single cell-RNA sequencing (scRNA-seq) experiments, AGM-EC from multiple derivations were thawed in parallel from low passage aliquots onto gelatin-treated T75 flasks in EC media. Confluent EC were cultured in serum-free medial (X-vivo 20) 24 hours prior to harvest by treatment with TrypLE Express, then re-suspended in PBS/10% FBS at 4°C, washed twice in PBS in 0.04% ultrapure BSA, and re-suspended PBS with 0.04% ultrapure BSA in on ice for scRNA-seq, as described below.

### AGM-EC co-culture

For co-culture experiments, AGM-EC at passage 15 or less were plated 24-48 hours prior to initiation of co-culture at a density of 1×10^4^ cells per well to gelatin-treated 96-well tissue culture plates for single cell index co-culture or 4×10^4^ cells per 24-well for bulk co-culture. For single cell index co-culture, AGM-derived haemogenic cells were individually index sorted to each well of 96-well containing AGM-EC in serum-free media consisting of X-vivo 20 with recombinant cytokines: stem cell factor (SCF) at 100 ng/ml, and interleukin-3 (IL-3) and thrombopoietin (TPO) each at 20 ng/ml. Following 5 to 7 days of co-culture, each well was visualized for haematopoietic colony formation and 50% of cells were harvested by vigorous pipetting for phenotypic analysis by flow cytometry, and in some experiments, remaining cells were used for confirmatory transplantation assay as previously described^26, 27^. For co-culture of bulk populations, sorted cells were re-suspended in serum-free culture media with cytokines, as above, and plated at 1-2 embryo equivalent of cells per 24-well containing AGM-EC. Following 6 to 7 days of co-culture, haematopoietic progeny were harvested by vigorous pipetting for subsequent analysis by flow cytometry and transplantation assays, as described below.

### Engineered culture conditions

Delta1ext-IgG (Dll1-Fc) was generated as previously described^72^. 48-well nontissue culture-treated plates (Falcon; BD Biosciences) were incubated with PBS containing: Dll1-Fc (2.5 μg/ml), anti-Notch1 (clone HMN1-12) and anti-Notch2 (clone HMN2-35) antibodies (each at 10 ug/ml), or Armenian Hamster IgG Isotype Control Antibody, together with 5 μg/ml recombinant fibronectin fragment (FN-CH-296), recombinant mouse VCAM-1/CD106 Fc Chimera, or human IgG control antibody, as indicated in results section. Following overnight incubation at 4°C, wells were washed twice with excess PBS prior to adding media. Serum-free media consisted of StemSpan SFEM II with recombinant cytokines: stem cell factor (SCF) at 100 ng/ml, and interleukin-3 (IL-3) and thrombopoietin (TPO) each at 20 ng/ml, and small molecule inhibitor of TGFBR (10 μM LY364947; from stock 10 mM solution in DMSO). For some experiments, as indicated, cells were cultured in above media with or without recombinant murine CXCL12 (100 ng/ml). Freshly sorted VE-cadherin^+^ECPR^+^CD61^+^ cells from dissected E9, E10 or E11 AGM (as indicated) were resuspended in media above and added to coated 48-well plates (approximately 1-2 embryo equivalent of cells per 48-well) in a total volume of 500 ul/48-well. Cells were harvested for phenotypic analysis by flow cytometry and transplantation assays at day 6-7 of culture, as described below.

### Flow cytometry analysis of cultured cells

Following co-culture, a fraction of the generated haematopoietic progeny was harvested by vigorous pipetting from the EC layer or engineered conditions for analysis of surface phenotype by flow cytometry (approximately 10% per well of cells cultured in bulk on AGM-EC or engineered conditions, or 50% of cells generated following single cell index culture on AGM-EC). Cells were spun and re-suspended in PBS with 2% FBS, pre-incubated with anti-mouse CD16/CD32 (FcRII block) and then stained with the following anti-mouse monoclonal antibodies: VE-Cadherin-PeCy7, CD45-PerCP, Gr-1-FITC, F4/80-FITC, Sca-1-APC, and EPCR-PE. DAPI was used to exclude dead cells. Flow cytometry was performed on a Becton Dickinson Canto 2 and data analyzed using FlowJo Software. Cells with HSC potential were identified phenotypically as VE-cadherin^-/low^CD45^+^Gr1^-^F4/80^-^ Sca1^high^EPCR^high^. We previously showed that detection of cells with this HSC phenotype following in vitro culture correlated with long-term multilineage engraftment as measured by transplantation assays performed in parallel, whereas haematopoietic progeny without detectable phenotypic HSC did not provide detectable long-term multilineage haematopoietic engraftment^26, 27^.

### Transplantation assays

Following co-culture, a fraction of the generated haematopoietic progeny was harvested by vigorous pipetting from the EC layer or engineered conditions, washed with PBS with 2% FBS and re-suspended in 100 ul PBS/2% FBS per mouse transplanted. For bulk co-culture experiments on AGM-EC or engineered conditions, the remaining 90% of cells in each well following flow cytometry analysis were pooled, washed with PBS with 2% FBS, and re-suspended at 1-2 embryo equivalent of cells per 100ul PBS/2% FBS, and combined with 5×10^4^ whole marrow cells from adult congenic C57BL/6.SJL-Ly5.1-Pep3b (CD45.1) mice in 100 ul PBS/2% FBS to provide haematopoietic rescue. For some single cell index assays, to validate the presence of functional HSC in colonies containing haematopoietic progeny with phenotypic HSC (VE-cadherin^-/low^CD45^+^Gr1^-^F4/80^-^Sca1^high^EPCR^high^) following co-culture, the remaining 50% volume of each 96-well was harvested for transplantation to individual mice, resuspended in 100ul, and combined with 5×10^4^ CD45.1 whole marrow cells in 100 ul PBS/2% FBS. Cell suspensions (200 ul total volume/mouse) were injected into lethally irradiated (1,000 cGy using a Cesium source) congenic CD45.1 adult recipients via the tail vein. For some experiments, secondary transplants were performed by transplanting 2×10^6^ whole bone marrow cells harvested from primary recipients to lethally irradiated CD45.1 secondary recipients via the tail vein. Flow cytometry analysis of peripheral blood obtained by retro-orbital bleeds was performed at 16 to 24 weeks following transplantation. Lineage-specific staining for donor (CD45.2) and recipient/rescue (CD45.1) cells from peripheral blood was performed as previously described^25^, using anti-mouse monoclonal antibodies indicated in Supplementary Table 7: CD45.1, CD45.2, CD3, CD19, Gr1, and F4/80. Multilineage engraftment was defined as >5% donor (CD45.2) contribution to the peripheral blood with contribution to each lineage of donor myeloid (Gr-1 and F4/80), B cells (CD19) and T-cells (CD3) detected at ≥0.5% at 16 to 24 weeks post-transplant, as indicated.

### Single cell RNA sequencing

For single cell RNA sequencing (scRNA-seq) studies, freshly sorted cells or cells harvested following culture (as indicated above) were washed with PBS containing 0.04% ultrapure BSA and re-suspended in 0.04% ultrapure BSA in PBS on ice. Cell suspensions were loaded into the Chromium Single Cell B Chip (10X Genomics) and processed in the Chromium single cell controller (10X Genomics), targeting 3500 cells per lane from freshly sorted AGM-derived cells, cultured AGM-EC, or progeny of AGM-derived cells following culture on AGM-EC or engineered conditions, or the remaining 50% progeny of clonal AGM-derived V^+^E^+^61^+^ following AGM-EC co-culture and flow cytometry analysis (estimated to contain < 500 cells). The 10X Genomics Version 2 single cell 3’ kit was used to prepare single cell mRNA libraries with the Chromium i7 Multiplex Kit, according to manufacturer protocols. Sequencing was performed for pooled libraries from each sample on an Illumina NextSeq 500 using the 75 cycle, high output kit, targeting a minimum of 50,000 reads per cell.

### Single cell transcriptome computational analysis

The Cell Ranger 2.1.1 pipeline (10X Genomics) was used to align reads to the mm10 reference genome and generate feature barcode matrix, filtering low-quality cells using default parameters. The Monocle (v.2.99.3) platform was used for downstream analysis, combining read-depth normalized data from replicate samples. Uniform Manifold Approximation (UMAP) was used for dimensionality reduction with the reduceDimension function. Clustering was performed by Louvain method with the clusterCells function. Differential gene expression between clusters (for Fig. 1) was performed using the principalGraphTest and find_cluster_markers functions, using a Moran’s I threshold of 0.25, filtering genes based on specificity >0.75. Gene ontology analysis was performed on genes identified as differentially expressed between clusters by PANTHER overrepresentation test with default settings (Fisher’s Exact, FDR correction) using annotation data sets for GO biological process and GO molecular function. Aggregated gene scores were calculated for each single cell as the log-transformed, size factor-normalized mean expression for each gene within a given GO term as indicated (see Supplementary Table 3). Cell type classification was performed using the classifyCells function (method set to “markers-only”), with marker gene sets based on established cell type-specific genes, as indicated. Pseudotime trajectory analysis was performed using the learnGraph function, with RGE_method set to SimplePPT. Differential gene expression over pseudotime was performed using the differentialGeneTest function, with fullModelFormulaStr set to Pseudotime, limited to genes with detected expression in >5% of cells and a significance cut-off of q-val < 0.01. Violin plots of normalized expression for each cell type were generated using the plot_genes_violin function. To identify genes coding for ligands expressed in niche endothelial cells, genes were first limited to those with detectable UMI in at least 10% of cells in 1) HSC-supportive AGM-EC #1, #2, and #3, (cluster 1, from Fig. 1) or 2) primary AGM-derived cells identified by cell type classification as “arterial EC” (from Fig. 3). These genes were then cross referenced with all potential ligands in the ligand-receptor database published by Ramilowski et al.^55^, using mouse genes corresponding to human genes in the published database. To identify genes coding for receptors expressed in developing HSC, genes were first limited to those with detectable UMI in 1) at least 10% of cells in primary AGM-derived cells identified by cell type classification as “HSC precursor” type, subclassified as “HE” type and “Pre-HSC” type (From Fig. 3), or 2) expressed in at least 20% of cells following AGM-EC co-culture identified by cell type classification as “HSC” type (from Fig. 4). These genes were then cross referenced with all potential receptors in the ligand-receptor database published by Ramilowski et al.^55^. All potential ligand-receptor pairings from this database were then determined for interactions between primary AGM-derived arterial EC or HSC-supportive AGM-EC stroma and primary AGM-derived HE/pre-HSC or in vitro generated HSC (Fig. 5 and Supplementary Table 6). Circos plots for visualization of ligand-receptor interactions for selected signal pathways (Fig. 5i-viii) were generated using the Circlize package^73^, modified from the iTALK package^74^. Ligands were categorized as those detected in either primary AGM-derived arterial EC, HSC-supportive AGM-EC lines, or both; or as those representing cytokines added exogenously during AGM-EC co-culture (ie. IL3, SCF, TPO) (Fig. 5 legend). Receptors were categorized as those detected in primary AGM-derived cells identified by cell type classification as “HSC precursor” type, subclassified as “HE” type and “Pre-HSC” type or following AGM-EC co-culture identified by cell type classification as “HSC” type (Fig. 5 legend). Receptors that were detected to significantly vary over pseudotime were also categorized as enriched in “HE” type vs “Pre-HSC” type (corresponding to early pseudotime, see Fig. 3E, Supplementary Table 4), enriched in “Pre-HSC vs HE” type (corresponding to late pseudotime, see Fig. 3E, Supplementary Table 4), or enriched in “HSC” type vs “HPC” type (corresponding to early pseudotime, see Fig. 4G, Supplementary Table 5). Ligands-receptor interactions involving ligands that were significantly enriched in expression in HSC-supportive versus non-supportive AGM-EC (from Supplementary Table 2) are indicated (Fig. 5 legend, yellow arrows). R scripts used for analysis are available upon request.

### Statistics

For statistical analysis between groups, two-tailed, unpaired Student’s t-test was used to calculate P values where indicated. P value less than 0.05 was considered significant.

### Data Availability

All sequencing data is available at NCBI GEO (Accession number GSE145886).

## ACKNOWLEDGEMENTS

Research reported in this publication was supported by the National Institute of Diabetes and Digestive and Kidney Diseases of the National Institutes of Health (NIH) under Award Number RC2DK114777 (to I.D.B., S.R., C.T., and B.K.H.) and National Heart, Lung, And Blood Institute of the NIH under Award Number K08HL140143 (to B.K.H.). B.K.H. is supported by the American Society of Hematology Scholar Award. We would like to thank David A. Flowers for superb technical assistance, the Fred Hutchinson Flow Cytometry Core for assistance with FACS, and members of the Trapnell, Hadland, Bernstein, and Rafii labs for constructive feedback.

## AUTHOR CONTRIBUTIONS

Conceptualization, B.K.H., I.D.B, S.R, J.B., and C.T.; Methodology, B.K.H., I.D.B., B.V., S.R, J.B., and C.T.; Investigation, B.K.H., B.V., T.D., S.D., D.J., and C.N. Writing – original draft, B.K.H.; Writing – Review & Editing, all authors; Funding acquisition, B.K.H., I.D.B., S.R., and C.T.; Supervision, B.K.H., I.D.B., S.R., and C.T.

## MATERIALS & CORRESPONDENCE

Correspondence and material requests should be addressed to Brandon Hadland at.

