## Supplemental Figures for "Engineering a niche supporting haematopoietic stem cell development using integrated single cell transcriptomics"

**Supplementary Tables and Figures**

**Supplementary Table 1.** Long-term peripheral blood engraftment data. (Related to Figs. 1B, 2B, 2C, 6A-C)

**Supplementary Table 2.** Cluster-specific genes differentially expressed in HSC-supportive AGM-EC (cluster 1) or non-supportive AGM-EC (Cluster 2), ordered by specificity. (Related to Fig. 1C-E).

**Supplementary Table 3.** Gene ontology terms (biological processes and molecular functions) associated with cluster-specific genes expressed in HSC-supportive AGM-EC. List of cluster-specific genes used for aggregated gene scores based on gene ontology terms. (Related to Fig. 1D).

**Supplementary Table 4.** Genes differentially expressed over pseudotime representing the EC/HE to haematopoietic transition, ordered by q-value. (Related to Fig. 3D-E).

**Supplementary Table 5.** Genes differentially expressed over pseudotime representing HSC to HPC differentiation, ordered by q-value. (Related to Fig. 4G).

**Supplementary Table 6.** List of ligand-receptor interactions between primary AGM-derived arterial EC or HSC-supportive AGM-EC stroma and primary AGM-derived HE/pre-HSC or in vitro generated HSC (Related to Fig. 5).

**Supplementary Table 7.** List antibodies and key reagents.

**Supplementary Figure 1**. **a**. Counts of UMI and unique genes expressed per cell for each AGM-EC sample (EC1-4). Boxplots show median values and interquartile ranges; upper/lower whiskers show 1.5X interquartile range with outliers shown as individual dots. **b**. Expression of cell cycle gene *Mki67,* demonstrating intra-cluster cell cycle heterogeneity. (Related to Fig. 1).

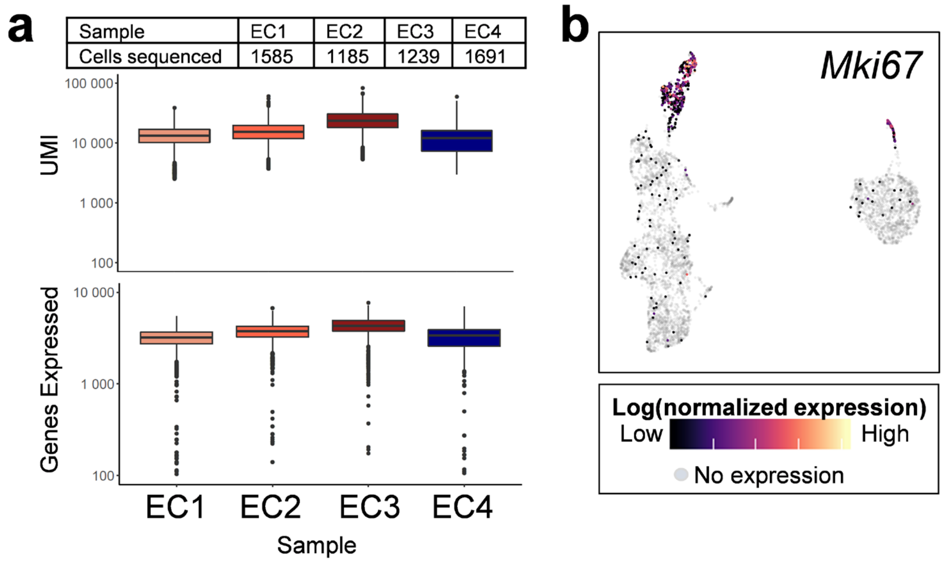

**Supplementary Figure 2**. Gating strategy with isotype controls for EPCR (IgG PerCP) and CD61 (IgG APC) within the population sorted as positive for VE-Cadherin. (Related to Fig. 2a).

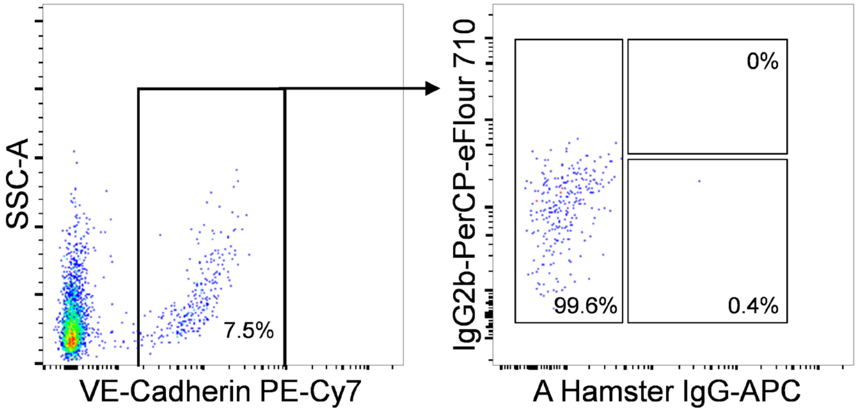

**Supplementary Figure 3. a-c**. Sort strategy (**a**) and post-sort analysis for E10 (**b**) and E11 (**c**) AGM V$^+$61$^+$E$^+$ cells isolated for scRNA-seq. **d**. Counts of UMI and unique genes expressed per cell for each embryo stage/sample. Boxplots show median values and interquartile ranges; upper/lower whiskers show 1.5X interquartile range with outliers shown as individual dots. **e**. UMAP with cells shown by embryo stage/sample. **f**. Genes used for cell type classification. **g**. Mature haematopoietic cell types identified in cluster 6. (Related to Fig. 3).

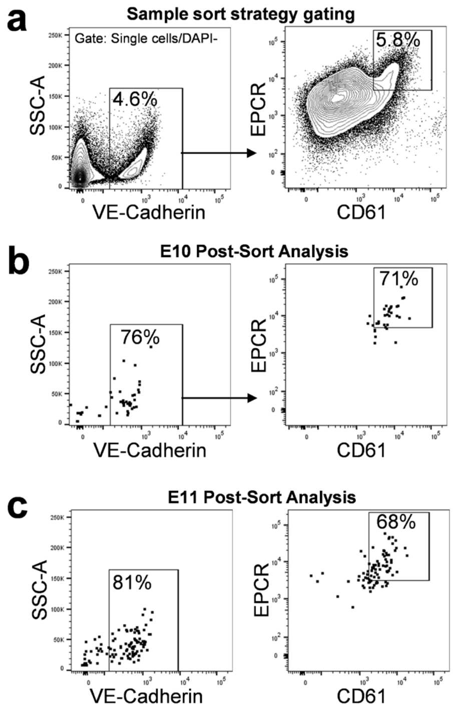


**d**

| Sample | E10 (31-39 sp) | E11 (40-45 sp) |
|---|---|---|
| Embryos pooled | 46 | 54 |
| Cells sequenced | 1622 | 1470 |

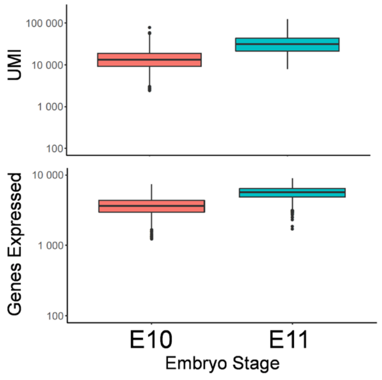


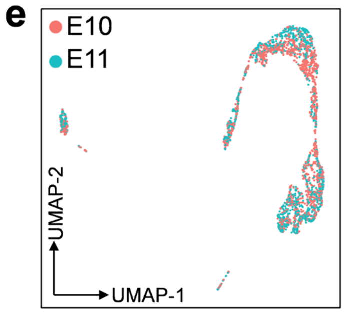


**f**

**From Figure 3C**

| Cell type | Genes expressed | Genes not expressed |
|---|---|---|
| Arterial EC | *Cdh5, Dll4, Efnb2, & Hey1* | *Runx1, Nr2f2, Nrp2* |
| Other EC (non-arterial) | *Cdh5, Nr2f2, & Nrp2* | *Runx1, Efnb2, Hey1, Meox1* |
| Hematopoietic | *Runx1 & Gfi1* | |
| Somite | *Pax1 & Meox1* or *Pax3 & Myf5* | |

**From Figure 3G-I**

| Cell type | Genes expressed | Genes not expressed |
|---|---|---|
| HSC Precursor | *Runx1, Pdzk1ip1, Ctnnal1* | *Flt3, Il7r, Fcgr3, Csf3r* |
| HE subtype | | *Ptprc, Itga2b* |
| Pre-HSC subtype | *Ptprc* or *Itga2b* | |
| Progenitor | *Runx1& Flt3, Il7r, Fcgr3,* or *Csf3r* | |

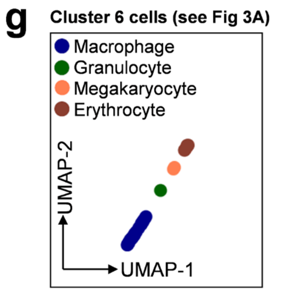


| Cell type | Genes expressed |
|---|---|
| Macrophage | *C1qb, Mrc1* |
| Granulocyte | *Fcnb* |
| Megakaryocyte | *Pf4, Gata1* |
| Erythrocyte | *Gypa, Klf1* |

**Supplementary Figure 4. a-b**. UMAP of single cell transcriptomes of the progeny of a single V+61+E+ haemogenic precursor following co-culture on AGM-EC, with (**a**) “HSC” phenotype and (**a**) mixed “HSC/HPC” phenotype, showing clusters classified by cell type as endothelial (EC) and haematopoietic (Hem), based on expression of VE-Cadherin (*Cdh5*) and CD45 (*Ptprc*) (Related to Fig. 4A-D). **c-d**. Counts of UMI per cell and unique genes expressed per cell for each cell type for samples shown in a-b, respectively. Boxplots show median values and interquartile ranges, upper/lower whiskers show 1.5X interquartile range, outliers shown as individual dots. **e**. Genes used for cell type classification of HSC and HPC. (Related to Fig. 4).

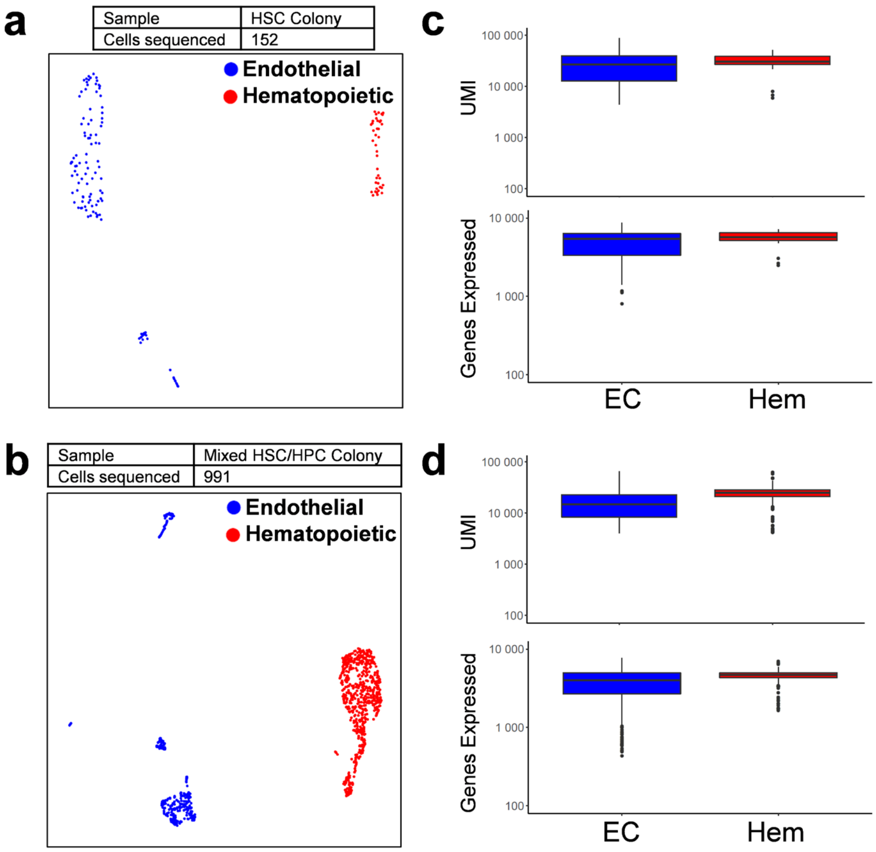


**e** **From Figure 4F**

| **Cell type** | **Genes expressed** | **Genes not expressed** |
|---|---|---|
| HSC Type | *Runx1, Pdzk1ip1, Vwf* | *Cd48, Itgal* |
| HPC Type | *Cd48* or *Itgal* | |

**Supplementary Figure 5. Single cell RNA-sequencing of AGM $V^+61^+E^+$ cells identifies transcriptional signatures of primary AGM niche arterial endothelial cells**. **a**. Comparison of surface expression of EPCR, CD61, CD41, and CD51 in VE-Cadherin$^+$ cells from primary AGM and cultured AGM-EC. **b**. Gene set scores representing top GO terms for transcripts differentially expressed by HSC-supportive verses non-supportive AGM-EC in single cell transcriptomes from primary AGM $V^+61^+E^+$ cells (see Fig. 1d). **c**. Gene expression heatmap in primary AGM $V^+61^+E^+$ cells for endothelial marker, *Kdr*, and transcripts differentially expressed between HSC-supportive and non-supportive AGM-EC (see Fig. 1e). (Related to Fig. 1, Fig. 3).

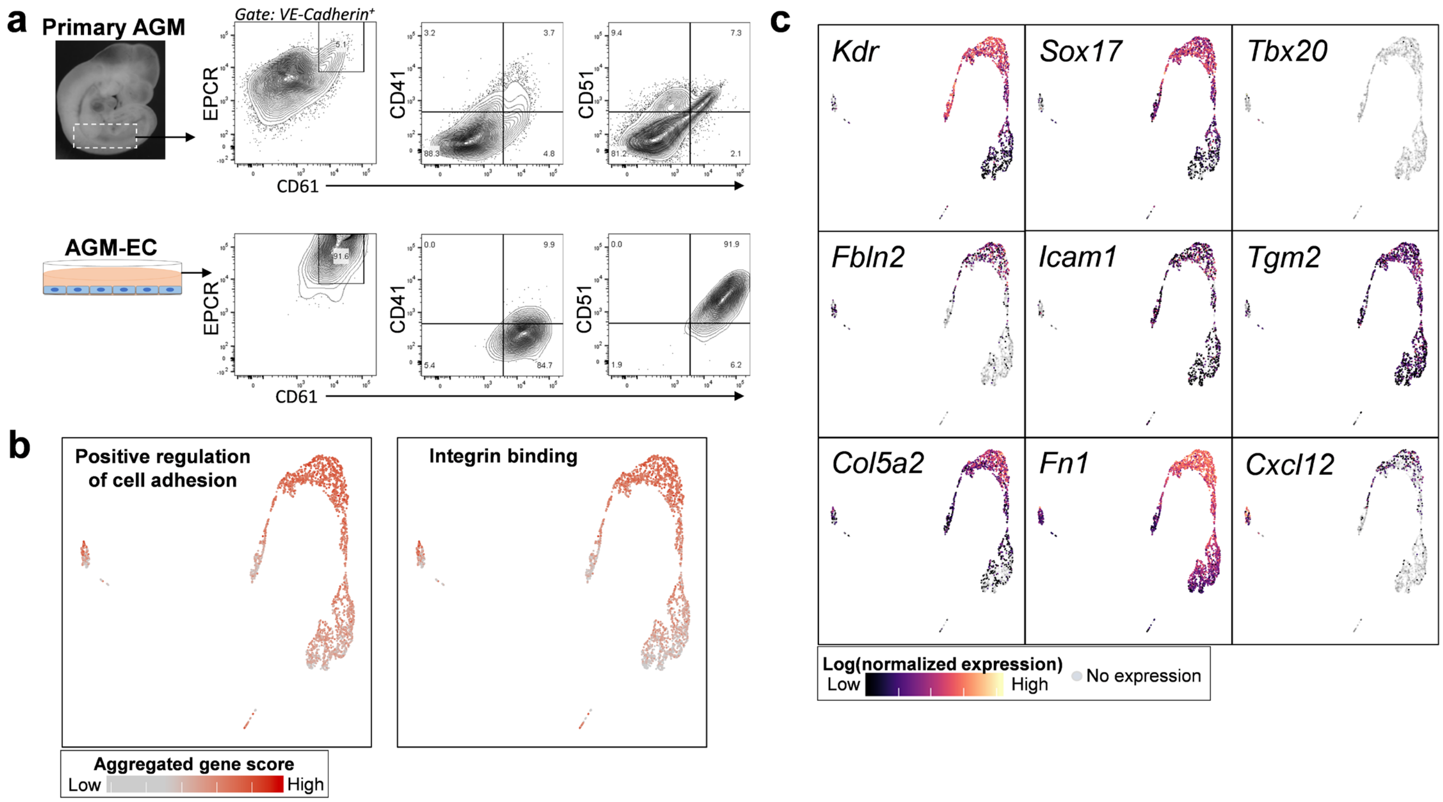

**Supplementary Figure 6.** A. Flow cytometric analysis of progeny of E11 AGM-derived $V^{+}61^{+}E^{+}$ cells follow culture in engineered conditions in the presence of immobilized Dll1-Fc, aN1/N2 Ab, or control (IgG), showing gating for phenotypic population of haematopoietic stem and progenitor cells (HSPC) (VE-$Cad^{low/-}CD45^{+}Gr1^{-}$ $F4/80^{-}Sca1^{+}EPCR^{+}$). B. Total live cells and C. Total HSPC generated per AGM equivalent of starting population following culture in engineered conditions in the presence of immobilized Dll1-Fc, aN1/N2 Ab, or control (IgG). (* $p < 0.001$) (Relate to Fig. 6B). D. Counts of UMI per cell and unique genes expressed per cell for each sample. Boxplots show median values and interquartile ranges, upper/lower whiskers show 1.5X interquartile range, outliers shown as individual dots. E. Genes for cell type classification. (Related to Fig. 6E-H).

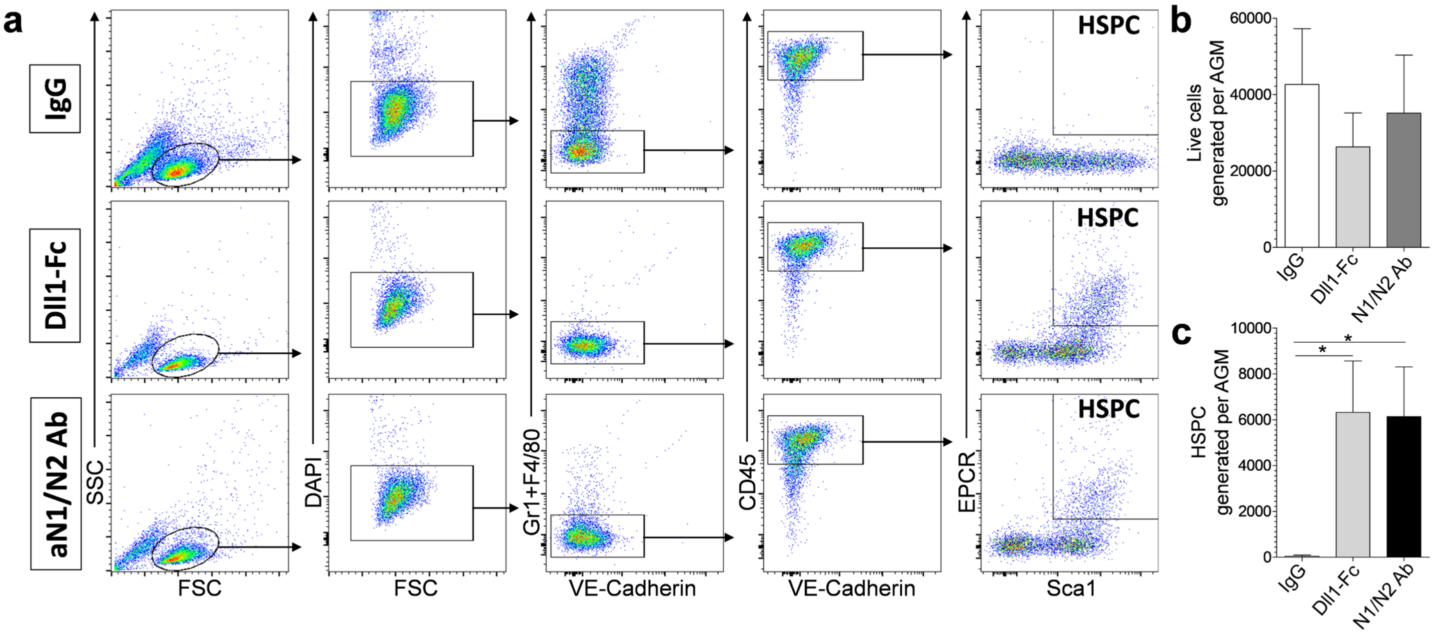


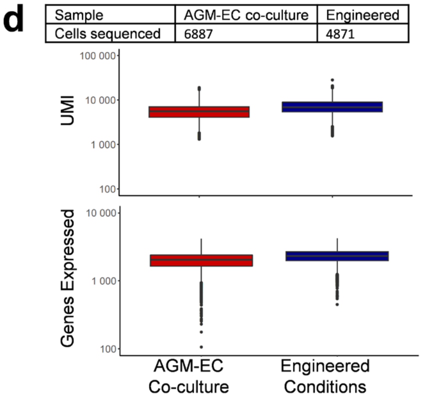


**e**

**From Figure 6F, 6H**

| **Cell type** | **Genes expressed** | **Genes not expressed** |
|---|---|---|
| HSC | *Pdzk1ip1, Hlf, Procr* | *Kdr* |
| MPP | *Kit, Cd34, Flt3* | *Pdzk1ip1* |
| T Prog | *Ptcra, Il7r* | |
| B Prog | *Ebf1, Il7r* | *Hlf, Pdzk1ip1* |
| Gran/Mono Prog | *Cd34, Fcgr3, & Cebpa, or Irf8 & Zeb2* | *Hlf, C1qb, Fcnb, Cd14, Gfi1* |
| Granulocyte | *Fcnb, Elane* | |
| Mono/Mac | *Cd14 & Adgre1, or C1qb & Cd36* | |
| Dendritic | *Siglech, Cd7* | |
| Mast | *Prss34, Cd200r3* | |
| Ery/Meg Prog | *Gata1, Gfi1b* | *Gypa, Cd34, Vwf, Prss34, Cd200r3* |
| Erythrocyte | *Gypa, Klf1* | |
| Megakaryocyte | *Vwf, Mpl* | *Hlf* |
| EC | *Cdh5, Kdr* | *Ptprc* |
